# Differential cellular responses to adhesive interactions with galectin-8 and fibronectin coated substrates

**DOI:** 10.1101/2020.06.14.127076

**Authors:** Wenhong Li, Ana Sancho, Jürgen Groll, Yehiel Zick, Alexander Bershadsky, Benjamin Geiger

**Affiliations:** Department of Immunology, Weizmann Institute of Science; Department of Functional Materials in Medicine and Dentistry and Bavarian Polymer Institute, University of Würzburg; Department of Automatic Control and Systems Engineering, University of the Basque Country UPV/EHU, San Sebastian, Spain; Department of Molecular Cell Biology, Weizmann Institute of Science; Mechanobiology Institute, National University of Singapore, Singapore

**Keywords:** actin cytoskeleton, AFM, cell spreading, extracellular matrix, focal adhesions, filopodia, myosin-II, Rho GTPases, SIM

## Abstract

The mechanisms underlying the cellular response to extracellular matrices (ECM), consisting of multiple adhesive ligands, each with distinct properties, are still poorly understood. Here we address this topic by monitoring the cellular responses to two very different extracellular adhesion molecules – fibronectin and galectin-8 – and to mixtures of the two. Fibronectin is one of the major integrin ligands, inducing cell spreading and development of focal adhesions associated with contractile stress fibers. Galectin-8 is a mammalian lectin, which specifically binds to β-galactoside residues present on some integrins, as well as to other cell surface receptors. We found marked differences in HeLa-JW cell spreading, assembly of focal adhesions and actomyosin stress fibers, and formation of adherent filopodia, on rigid flat substrates functionalized by fibronectin or galectin-8 alone, or by mixtures of these two proteins. Spreading on galectin-8 resulted in a larger projected cell area compared to that on fibronectin, by more extensive formation of filopodia, coupled with an inability to activate focal adhesion and stress fiber assembly. These differences could be partially reversed by experimental manipulations of small G-proteins of the Rho family and their downstream targets, such as formins, the Arp2/3 complex, and Rho kinase. Another factor affecting the spreading process was shown to be the enhanced physical adhesion of the cells to galectin-8, as compared to fibronectin. Notably, at least one process, the formation of adherent filopodia, was synergistically upregulated by both ligands, so filopodia development on the substrate coated with a mixture of fibronectin and galectin-8 was far more prominent than on each ligand alone.

## Introduction

In multicellular organisms, the majority of cells interact with the extracellular matrix (ECM), a multimolecular network comprising the cells’ microenvironment (Naba et al., 2016). ECM components are synthesized, secreted and assembled by cells, and form specialized tissue scaffolds characterized by specific biochemical, topographical and mechanical features. ECM networks serve as primary sources of environmental information for many cell types, normal and transformed, affecting their shape, adhesive properties, cytoskeletal organization, and migration (Bonnans et al., 2014; Humphrey et al., 2014; Muncie and Weaver, 2018).

Several classes of cell surface receptors interact with the ECM, and convey to cells its biochemical and mechanical characteristics. Among these are transmembrane receptors of the integrin family (24 types in mammalian cells), and transmembrane proteoglycans such as syndecans, CD44, betaglycan, and neuropilin-1, each of which generate different types of signals (Humphries et al., 2019; Multhaupt et al., 2016). Depending on the type of matrix, its mechanical characteristics, as well as the cellular context, transmembrane matrix receptors assemble into different types of adhesion complexes via which they interact with specific cytoskeletal structures. Adhesion complexes such as focal adhesions (Geiger et al., 2009), podosomes (Alonso et al., 2019; Schachtner et al., 2013), hemidesmosomes (Walko et al., 2015), filopodia (Jacquemet et al., 2015) and adhesion waves (Case and Waterman, 2011) are formed by several hundreds of structural and signaling proteins, which collectively mediate the force-transducing, mechanosensory, and signal-generating functions of these structures.

Thus far, our knowledge and understanding of the processes underlying matrix-dependent signaling, formation of adhesion complexes, and downstream cellular responses, are based on studies of a limited number of distinct matrix components. In most of these studies, specific matrix proteins such as fibronectin, collagen or laminins were used as 2-dimensional adhesive substrates. In more recent studies, these proteins were presented to cells on substrates of variable topographies and rigidities, as well as in the form of 3-dimensional networks. The latter studies revealed different types of cellular responses, depending on the mechanical and topographical characteristics of the substrates.

At the same time, it became clear that even under similar topographical and mechanical conditions, different types of matrix proteins can generate different and sometimes contradictory cellular responses. For example, while adhesion of fibroblasts to a planar rigid substrate coated with a mixture of fibrin and fibronectin triggers RhoA activation and subsequent formation of stress fibers and focal adhesions, addition of another matrix protein, tenascin-C, a ligand for a specific subset of integrins (Midwood et al., 2016), strongly suppresses RhoA activation and focal adhesion formation (Midwood and Schwarzbauer, 2002; Wenk et al., 2000). Another major matrix protein, thrombospondin-1, activates Rac1 and Cdc42 more strongly than fibronectin, resulting in the activation of PAK kinase and the extension of numerous fascin-containing filopodia (Adams and Schwartz, 2000). The cell matrix receptor syndecan-1 was thought to be responsible for these thrombospondin-1 effects (Adams et al., 2001), even though thrombospondin-1 also binds several other types of receptors including integrins (Resovi et al., 2014). For the majority of matrix-associated proteins, however, not only are downstream signaling pathways insufficiently understood, but even the basic phenotypic effects are poorly characterized.

Galectins comprise a large family of animal lectins, expressed in multiple tissues and displaying diverse functions, including the regulation of cell adhesion and migration. They are secreted by cells via non-conventional mechanisms bypassing the endoplasmic reticulum/Golgi dependent secretary pathway (Popa et al., 2018), and function as matricellular proteins serving as soluble ligands cross-linking molecules on the cell surface, as well as components of the extracellular matrix (Elola et al., 2007; He and Baum, 2006; Nabi et al., 2015). All galectins bind β-galactoside and interact with membrane glycoproteins and glycolipids. Depending on the organization of sugar moieties and protein structures, different galectins can interact with specific molecules on the cell surface. Galectins are expressed in all tissues and display multiple functions, including in particular the regulation of cell adhesion and migration.

A prominent member of the galectin family is galectin-8, a molecule containing two carbohydrate-recognition domains (CRDs), connected by linkers of variable lengths in three different isoforms (Gal-8S, Gal-8M, Gal-8L). Galectin-8 plays an important role in normal physiological processes such as vascular and lymphatic angiogenesis (Troncoso et al., 2014), platelet activation (Romaniuk et al., 2015; Romaniuk et al., 2010), cell spreading, polarization of T-lymphocytes (Cárcamo et al., 2006), and limb development (Newman et al., 2018). At the same time, galectin-8 is often overexpressed and secreted by some types of tumor cells (Elola et al., 2014; Vinik et al., 2018), a phenomenon thought to be critical to their metastatic ability (Gentilini et al., 2017). The common denominator underlying galectin-8 involvement in these diverse processes might be its ability to regulate cell adhesion (Zick et al., 2002).

Galectin-8 immobilized on a rigid substrate can support the adhesion and spreading of different cell types, e.g. T-cells, trabecular meshwork cells, normal and neoplastically transformed cultured fibroblast and epithelial cells of many types (Cárcamo et al., 2006; Diskin et al., 2012; Levy et al., 2001), and initiate downstream adhesion-dependent signaling such as tyrosine phosphorylation of focal adhesion kinases (FAK) and paxillin (Diskin et al., 2012; Levy et al., 2001). There also exists evidence that the small G-proteins Rho and Rac1 are activated upon cell attachment to a galectin-8 coated substrate (Cárcamo et al., 2006; Diskin et al., 2012).

Both N-terminal and C-terminal carbohydrate recognition domains (CRDs) of galectin-8, as well as the linker connecting them, are required for optimal cell adhesion (Levy et al., 2006). During adhesion, cells interact with galectin-8 via a subset of glycosylated integrins such as α3, α6, and ß1 in human lung carcinoma 1299 cells; α1ß1, α3ß1, and α5ß1 in Jurkat T cells; and α3ß1, α5ß1, and αvβ1 in trabecular meshwork cells (árcamo et al., 2006; Diskin et al., 2009; Elola et al., 2014; Hadari et al., 2000). In addition, cells can interact with galectin-8 via other classes of receptors; for example, ALCAM/CD166 in vascular endothelial cells (Troncoso et al., 2014) and breast cancer cells (Fernández et al., 2016; Ferragut et al., 2019); podoplanin in lymphatic endothelial cells (Cueni and Detmar, 2009; Troncoso et al., 2014) and in cancer-associated macrophages (Bieniasz-Krzywiec et al., 2019); and urokinase plasminogen-activated receptor (uPAR) in osteoblasts (Vinik et al., 2015).

Information about the cellular response to contact with galectin-8 that underlies cell spreading behavior is, however, sparse. The processes of actomyosin cytoskeleton organization, formation of lamellipodial and filopodial protrusions, and assembly of adhesion structures upon cell plating on a galectin-8 coated substrate, have as yet been insufficiently studied. Even less is known about cell reactions to composite substrates containing galectin-8, together with other matrix proteins. In particular, galectin-8 can interact with fibronectin (Levy et al., 2001; Reticker-Flynn et al., 2012), and some data suggest that substrates containing a mixture of these proteins enhance the spreading of metastatic cell (Reticker-Flynn et al., 2012). However, the changes in cell spreading reactions underlying such behavior remain uncharacterized.

In this study, we investigated, in detail the process of cell spreading on galectin-8 coated substrates, in comparison to such spreading on fibronectin. We found marked differences in self-organization of the actomyosin cytoskeleton, formation of lamellipodial and filopodial protrusions, and assembly of adhesion contacts upon cell adhesion to these two types of matrix proteins. These differences depended on much stronger adhesion forces, as well as on alterations in the balance between activities of small Rho family proteins and their downstream targets when cells attached to galectin-8, as compared to fibronectin. We further investigated the spreading of cells on a composite substrate formed by mixing of galectin-8 and fibronectin at different ratios, and found strong synergy between fibronectin and galectin-8 underlying the formation of adhesive filopodial protrusions. These observations could help to explain the apparent involvement of galectin-8 in establishing cancer cell phenotypes.

## Materials and Methods

### Cell culture, DNA constructs, and reagents

Hela-JW is a subline of a HeLa cervical carcinoma cell line derived in the laboratory of J. Wiliams (Carnegie-Mellon University, USA) on the basis of better attachment to plastic dishes (Bai et al., 1993; Paran et al., 2006). The cells were cultured in Dulbecco’s modified Eagle’s medium (DMEM) supplemented with 10% fetal bovine serum (FBS), 1 mM sodium pyruvate and 100 U/mL penicillin-streptomycin in 5% CO2 incubator at 37 °C. The cell culture reagents were purchased from Biological Industries, Ltd. (Beit Haemek, Israel), and used according to the manufacturer’s instructions, unless otherwise stated. No cell lines used in this study were found in the database of commonly misidentified cell lines that is maintained by ICLAC and NCBI Biosample. We did not attempt to authenticate them.

YFP-tagged paxillin construct in pEYFP vector (Zaidel-Bar et al., 2007) was used to derive Hela-JW subline stably expressing paxillin which was kindly provided to us by Dr. S. W. Katz. The cells were also transiently transfected with the following DNA plasmids: tdTomato-F-tractin (Schell et al., 2001) (a gift from M. J. Schell, Uniformed Services University, Bethesda, Maryland), myosin II regulatory light chain MRLC-GFP (Kengyel et al., 2010) (a gift from Drs. W. Wolf and R. Chisholm, Northwestern University, Chicago, Illinois, USA), mDia2 ΔDAD-GFP cloned by Dr. N. O. Alieva in Bershadsky’s lab (Alieva et al., 2019), FMNL2 ΔDAD-GFP cloned in Dr. E. Manser’s lab (A-star, Singapore). All the transfections were done using Lipofectamin 2000 (Invitrogen^™^) following the manufacturer’s protocols.

### Transfection of siRNA

Cells were seeded into a 35 mm dish on day 0 and transfected with 20 μM of Rac1, Cdc42, FMNL2, mDia2, and Arp2 siRNA (Dharmacon, ON-TARGET plus SMART pool siRNA, catalogue no. L-011195-00-0005) using Lipofectamine RNAiMAX (Invitrogen) on day 1 and day 2. Control cells were transfected with scrambled control siRNA (Dharmacon, ON-TARGET plus Non-targeting pool siRNA, catalogue no. D-001810-10). Cells were imaged on day 4. The four siRNA sequences in the smart pool were as follows. siCDC42: GGAGAACCAUAUACUCUUG, GAUUACGACCGCUGAGUUA, GAUGACCCCUCUACUAUUG, CGGAAUAUGUACCGACUGU, siRac1: UAAGGAGAUUGGUGCUGUA, UAAAGACACGAUCGAGAAA, CGGCACCACUGUCCCAACA, AUGAAAGUGUCACGGGUAA, siFMNL2: GAACCUACCUCCUGACAAA, UAAGAGAACUGGAAAUUUC, UAACAGACAUGUAUAUGAG, AAUUAGGCCUGGACGAAUA, siDIAPH3: GAUCAGACCUCAUGAAAUG, GAGAAGAAAUCGAUUAAGA, GUAUGCAGCUCAUCAAUGC, GUAGACAUUUGCAUAGAUC, siArp2: GAAGUUAACUACCCUAUGG, GCAAGUGAAUUACGAUCAA, GAAACGGUUCGCAUGAUUA, UGGUGUGACUGUUCGAUAA

### Substrate coating

Bacterially expressed recombinant galectin-8 were purified as previously described (Hadari et al., 1995). α-Lactose-Agarose beads used for galectin-8 purification were purchased from Sigma (Catalog no. L7634). Galectin-8 mutated forms were generated as previously described (Levy et al., 2006). For some experiments, galectin-8 was labeled by Alexa Fluor 568 Protein Labeling Kit (Molecular Probes, Thermo Fisher Scientific) according to the manufacturer’s instructions. Fibronectin Solution (Bovine) at 1mg/ml was purchased from Biological Industries (03-090-1-01). Fibronectin HiLyte488^™^ was purchased from ENDO scientific services, Israel (Catalog no. FNR02-A).

The glass-bottomed petri-dish (MatTek, P35G-1.5-14-C) was coated with 25 μg/ml galectin-8 or fibronectin solution in PBS or by their mixture prepared by gentle pipetting in an Eppendorf microtube (1.5 ml). The dish was incubated with protein solution for two hours at room temperature and washed five times with PBS. In special control experiments with fluorescently labeled fibronectin and galectin-8, we checked that the concentration of both proteins that we used were saturating so that its absorption on the glass was maximal. We chose 25 μg/ml because the absorption of the proteins onto the cover glass reaches plateau for both fibronectin and galectin-8 at this concentration. Even though the presence of 25 μg/ml of galectin-8 somewhat reduced the absorption of fibronectin, at the 25 μg/ml of fibronectin, such reduction was minimal (Supplementary Figure 8). The cells were seeded onto the protein coated cover glass in DMEM medium without serum.

### Cell suspension preparation

To study the cell spreading on fibronectin and galectin-8 coated substrate, the cells from 70%-80% confluent culture in 22.1 mm well of multi-well dish were first washed with warm PBS once, then incubated in 150 μl of Trypsin EDTA Solution B (Trysin 0.25%, EDTA 0.05%) (Biological Industries USA, 03-052-1B) at 37°C for two minutes and gently suspended by addition of 5 ml serum-free DMEM with trypsin inhibitor (T9003, Sigma-Aldrich) (1mg of trypsin inhibitor per milliliter Trypsin EDTA Solution B). The suspension was centrifuged at 1000 rpm for 5 minutes, supernatant was removed and serum-free medium was added to re-suspend the cells. Then, the cells were plated to the pre-coated petri dish and either imaged or fixed at appropriate time points.

### Drug treatment

The Rho activator, CN03 (Cytoskeleton Inc. USA, catalogue number: CN03), was added at concentration of 1 μM 30 minutes following cell plating onto fibronectin or galectin-8 coated substrates, and incubated 3 hours more before fixation and staining. For ROCK kinase inhibition studies, the suspended cells were pretreated with 100 μM Y27632 in DMEM serum free medium for half an hour at 37°C and then were allowed to attach to either fibronectin or galectin-8 coated substrate in the presence of the inhibitor. For sugar inhibition studies, 10 mM thiodigalactoside (catalogue no. 17154, Cayman Chemical) was added to cells in suspension and incubated for 10 minutes at 37°C before the cells were seeded on the substrates. The RGD peptide (CGGGRGD, GeneCust, catalog number: HY-P2219), at a final working concentration of 10 μg/ml, was also added to cell suspension 10 minutes before plating and then remained in the medium during the experiments.

### Immunoblotting and small GTPases pull down assay

Transfected cells were lysed in RIPA buffer on day 4 and proteins extracted were separated by 10% - 15% SDS polyacrylamide gel and transferred to Nitrocellulose membranes (Bio-Rad) at 100V for 1 hour and blocked for 1 hour with 5% non-fat milk (Bio-Rad) before incubation at 4 C overnight with appropriate primary antibodies: Rho Rabbit Antibody (Cell Signaling, catalogue no. 8820), Rac1 Mouse Antibody (Cell Signaling, catalogue no. 8815), Cdc42 Mouse mAb (Cell Signaling, catalogue no. 8819), Anti--tubulin (Sigma, catalogue no. T6199, dilution 1:5,000, clone DM1A) was used as an internal control. The primary antibody binding was processed for ECL detection (Bio-Rad) with appropriate HRP-conjugated secondary antibodies (Santa Cruz Biotechnology, catalogue no. sc-2004/5, dilution 1:10,000).

Hela-JW cells were plated on fibronectin or galectin-8 coated substrates, and lysed half an hour after plating, and the active form of RhoA, Rac1, and Cdc42 were extracted using Active Rho Detection Kit from Cell Signalling (Catalog. 8815, 8819, 8820) following the manufacture’s protocol.

### Cell Adhesion Forces

Two defined regions of the glass bottom petri dish (GWSB-5030, WillCo Wells) had been freshly coated beforehand with galectin and fibronectin respectively, as explained above. Adhesion forces of cells to the underlying substrate for a contact time of 5 minutes were measured by Single Cell Force Spectroscopy (SCFS) using the FluidFM® technology (Cytosurge AG, Switzerland) incorporated to an Atomic Force Microscope (AFM) Flex-FPM system (Nanosurf GmbH, Germany) (Guillaume-Gentil et al., 2014). The system was mounted on an Axio Observer Z1 inverted microscope (Carl Zeiss, Germany) for the visualization of the cells. Micropipette cantilevers (Cytosurge, Switzerland) with an aperture of 4 μm in diameter and 0.3 N/m nominal spring constant were used. A cell was immobilized at the tip of the cantilever by applying a soft underpresure, then, it was brought into contact with the corresponding substrate by approaching at a speed of 1 μm/s until a set point of 5 nN was reached. This force was kept constant during the 5 minutes that the cell was kept in contact with the test material. After this time, the cantilever holding the cell was retracted from the surface, and its deflection during the retraction was recorded (Sancho et al., 2017). The deflection of the cantilever is directly proportional to the force exerted by the cells against the substrate while they are being pulled away from it and the maximum force peak is used as the indicator of cell adhesion force (Potthoff et al., 2012). Ten individual cells were measured under each experimental condition.

### Immunofluorescence staining

For immunostaining, cells cultured on glass bottomed dish were fixed/permeabilized in phosphate-buffered saline (PBS) containing 0.25% Triton X-100, 0.25% glutaradehyde and 3% paraformaldehyde at 37C for 15 min. The cells were then washed twice with PBS for 10 min each time. Before staining, the fixed cells were treated with 1mg/ml sodium borohydride in PBS for 15 min on ice. The cells were then washed with PBS, incubated with blocking solution (5% bovine serum albumin (BSA) in PBS) for 1 h at room temperature and washed with PBS again. The cells were incubated with appropriate primary antibodies (anti-paxillin, anti-myosin IIA, and anti-vinculin) at room temperature for one hour, and after triple 10 minutes washing with PBS, with appropriate fluorescently labelled secondary antibody and phalloidin to visualize actin. Goat anti-Rabbit/mouse IgG (H+L) Cross-Adsorbed ReadyProbes™ Secondary Antibody, Alexa Fluor 488/647 was purchased from Thermo Fisher (catalogue no. R37116/ R37114, A21245/A32728), and was used at dilution 1:400. Phalloidin–Tetramethylrhodamine B isothiocyanate was obtained from Sigma-Aldrich (catalogue no. P1951) and was used at 1:400 dilution. Purified Mouse Anti-Paxillin (BD Transduction Laboratories™, Clone: 349, catalogue no. 610052) was used at dilution 1:200, anti-myosin IIA, non-muscle antibody produced in rabbit (Sigma, catalogue no. M8064) at dilution 1:400. Monoclonal human vinculin antibody prepared by the Antibody Production Laboratory of the Department of Biological Services, Weizmann Institute of Science: (http://bioservices.weizmann.ac.il/antibody/about.html) was used at 1:50 dilution.

### Microscopy and live cell imaging

Cells were plated at a density of 5 × 10^4^ cells ml^-1^ onto the 35 mm cell culture dish with 14 mm-diameter glass bottom (MatTek, catalogue number: P35G-1.5-14-C) coated with fibronectin, galectin-8 or their mixture as described above. Cells were imaged in a medium with low level of background fluorescence FluoroBrite DMEM (Thermo Fisher, catalogue number: A1896701). Video recordings started 5 minutes after the cells were added to the dish. Differential interference contrast (DIC) and interference reflection microscopy (IRM) time-lapse imaging were carried out using the DeltaVision RT microscopy system (Applied Precision Inc., Issaquah, WA, USA), equipped with a ×60 oil immersion objective (1.40 NA, UPlanSApo), at 2 or 10 seconds time intervals between frames. Total internal reflection fluorescent (TIRF) images were acquired using the DeltaVision Elite microscopy system equipped with a multi-line TIRF module (Applied Precision, Inc.), and time-lapse movies were taken at 30-seconds intervals, unless otherwise indicated. Super-resolution SIM imaging was performed using W1-spinning-disc confocal unit coupled with the live super-resolution (SR) module (spinning disk based structured illumination super resolution (York et al., 2013)) (GatacaSystems), mounted on Eclipse microscope with Perfect Focus System, supplemented with the objective Plan Apo 100x oil NA1.45 and scientific complementary metal–oxide–semiconductor (sCMOS) camera Prime95B (Photometrics). Laser lines wavelength 488, 561nm were used.

### Image analysis

Projected cell area measurements: IRM images of cells taken at different time intervals after plating were first subtracted by a background image taken before plating the cell. Intensity thresholding by Otsu’s method (Otsu, 1979) was then applied to segment the cell. After the segmentation, individual objects were identified, and the object with the largest area was preserved and considered as the main cell body. For the measurement of projected cell area of fixed cells, we applied the same algorithm to actin fluorescence images instead of IRM images.

Filopodia measurements. To identify the filopodia and quantify their length and number in IRM and actin fluorescence images, we used FiloDetect algorithm (Nilufar et al., 2013). Filopodia were defined as high aspect ratio (≥1.5:1) objects protruding from the “main cell body” with a smooth boundary. The filopodia shorter than 0.6 μm were ignored.

Myosin II filament measurements. The live-cell recording of myosin II were performed by SIM imaging. A thresholding algorithm was first applied to actin images to segment the cell from the background. Then the cell were segmented into 1-pixel-width rings with the same contour as the cell edge without filopodia. The average GFP-MRLC intensity was calculated for each ring.

Paxillin structure measurements. Ilastik software(Berg et al., 2019) was applied to the paxillin images. The software runs a random forest classifier(Antonio et al., 2012) to the images to segment paxillin from the background. After the segmentation with the Ilastik software, the fluorescent intensity and area were analyzed using Matlab.

## Results

### The differential dynamics of filopodia and lamellipodia extensions on fibronectin and galectin-8

HeLa cell spreading manifests itself primarily in the continuous extension of two types of membrane protrusions, filopodia and lamellipodia. We used time-lapse interference reflection microscopy (IRM), along with differential interference contrast (DIC) microscopy, to visualize these processes (Fig. 1a, b; Movies 1, 2, 3). On fibronectin, the spreading process was initiated by the formation of filopodia (Fig. 1a; Movie 1). The filopodia were succeeded by blebs and irregular lamellipodial protrusions, which extended, paused or sometimes retracted (Fig. 1a; Movie 1). Usually, the lamellipodial spreading consisted of short periods of rapid protrusion, alternating with prolonged pause periods (Fig. 1c), so that the net rate of protrusion at the cell’s edge was slower than the rate of extension of individual lamellipodia (Fig. 1c; Movie 1).

**Figure 1:**
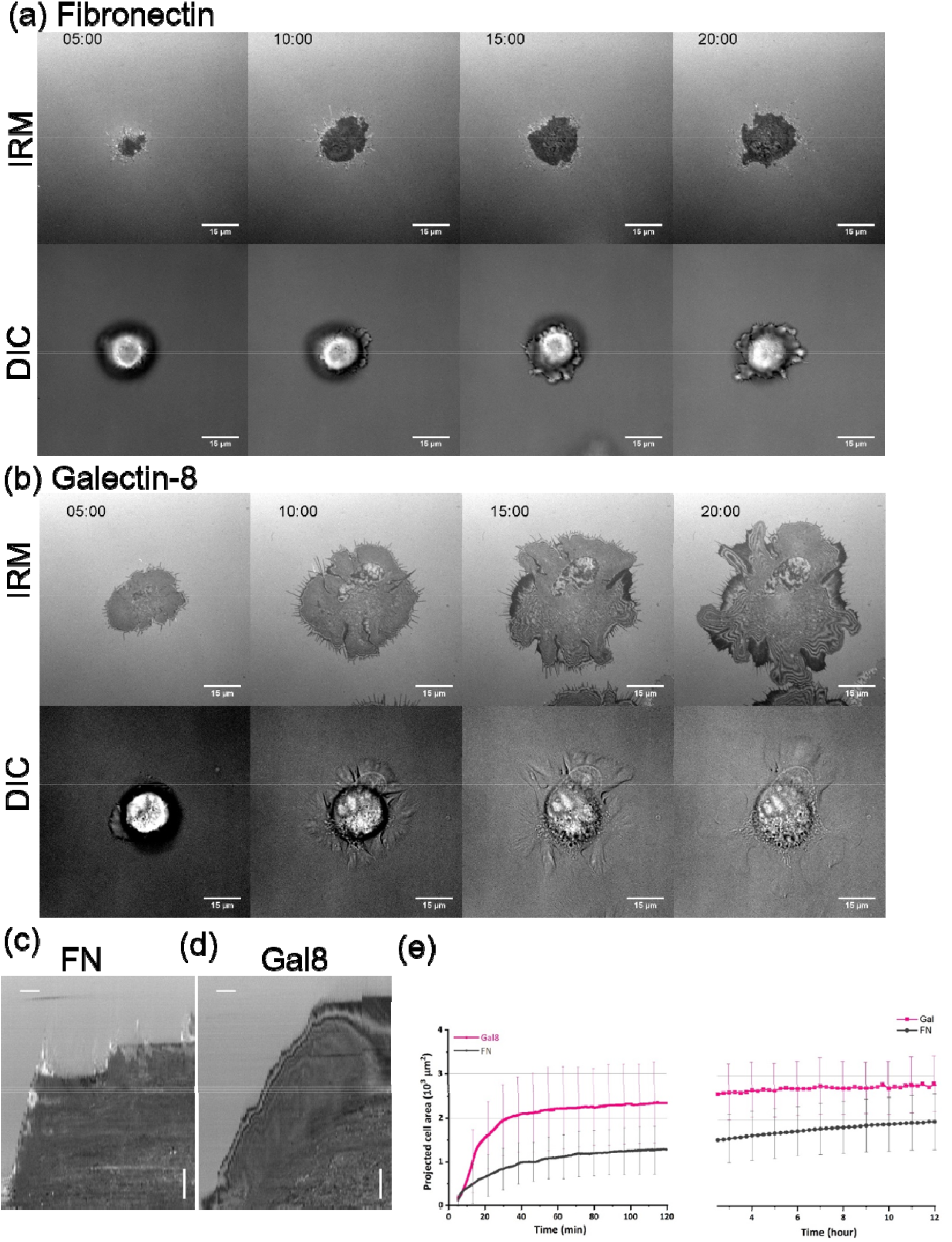
Spreading of HelaJW cells on fibronectin and galectin-8 coated substrates. (a) and (b) Time course of cell spreading on fibronectin (a) and galectin-8 coated substrates (b) imaged by interference reflection microscopy (IRM) (first rows) and differential interference contrast microscopy (DIC) (second rows) at different time intervals (minutes) (see also Movie 1 and Movie 2). Scale bars: 15 μm. Note that the projected cell area on galectin-8 dramatically exceeds that on fibronectin, due to faster extension of lamella. Numerous filopodia are seen in DIC images of cells on galectin-8. (c) and (d) Typical kymographs depicting the process of cell spreading on fibronectin (c) and galectin-8 (d). Images of a one pixel-wide strip perpendicular to the cell edge, taken every 10 seconds, are mounted along the horizontal time axis. The horizontal scale bar is 100 seconds, and the vertical scale bar is 2 μm. (e) Quantification of projected cell area on fibronectin and galectin-8 at different time points after cell plating. Error bar indicates standard deviation. Six cells for each condition were included in the spreading quantification for the spreading before 2 hours (left), and 20 cells for each condition were included in the quantification for the spreading after 2.5 hours (right).

On galectin-8, cell spreading was also initiated by the formation of filopodia (Fig. 1b, Fig. 2a; Movies 2, 3), which were rapidly succeeded by formation of lamellipodia. The rate of lamellipodial extension was somewhat slower than that seen on fibronectin (Fig. 1c, d). However, the extension of lamellipodia proceeded continuously without pause (Fig. 1b, d; Movies 2, 3). As a result, the net increase of projected cell area on galectin-8 was considerably more rapid, compared to that seen on fibronectin (Fig. 1e). When the velocity of lamellipodia spreading slowed, filopodial growth increased in speed, and new filopodia extending from the lamellipodial edge often formed (Fig. 2b). Similarly, filopodial growth cessation was accompanied by an increase in the rate of lamellipodial extension. Thus, on galectin-8, the spreading was not interrupted by pauses and proceeded via alternating waves of extension of lamellipodia and filopodia (Fig. 2b; Movies 2, 3). Another peculiar feature of cell spreading on galectin-8 was the “petaloid” shape of the cell contour. Extension of lamellipodia and filopodia occurred independently in 3-8 segments (“petals”) of the cell periphery (Fig. 2a; Movies 2, 3). This petaloid shape was characteristic of early stages of spreading. Later, the neighboring “petals” could fuse, and the spreading became more isotropic.

**Figure 2:**
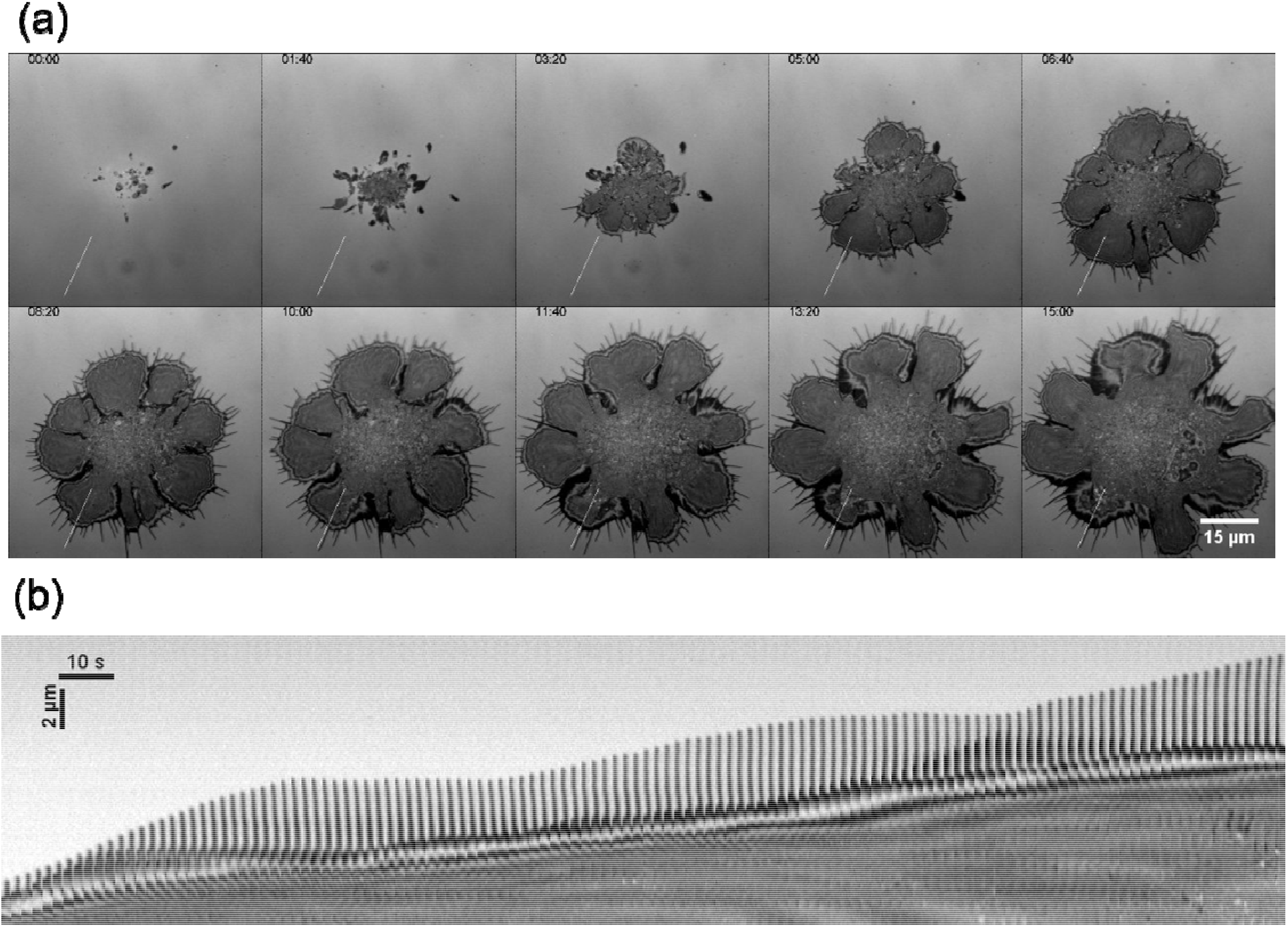
Filopodia and lamellipodia dynamics during cell spreading on galectin-8. (a) IRM images of a cell spreading on galectin-8 at different time intervals after plating (minutes) show petaloid lamellar extensions and filopodia. (b) Kymograph showing the time course of filopodia and lamellipodia protrusions from the strip with 1 μm width and 16 μm length crossing the cell periphery perpendicularly to the lamellipodia front, as indicated in (a). Note alternating waves of filopodia and lamellipodia extensions.

### The actomyosin cytoskeleton organizes itself differently in cells spreading on fibronectin and galectin-8

In agreement with previous studies (Gauthier et al., 2012; Wolfenson et al., 2014), we showed that spreading on fibronectin proceeded via formation of actin-rich lamellipodia and filopodia, followed by assembly of the actomyosin stress fiber system (Fig. 3a; Movie 4). The myosin II filaments visualized by cell transfection with GFP-myosin light chain appeared at the cell periphery, first moving centripetally (Hu et al., 2017), but later concentrating in large stress fiber-type actin bundles delineating the edges of polygonal cells (Fig. 3a; Movie 4).

**Figure 3:**
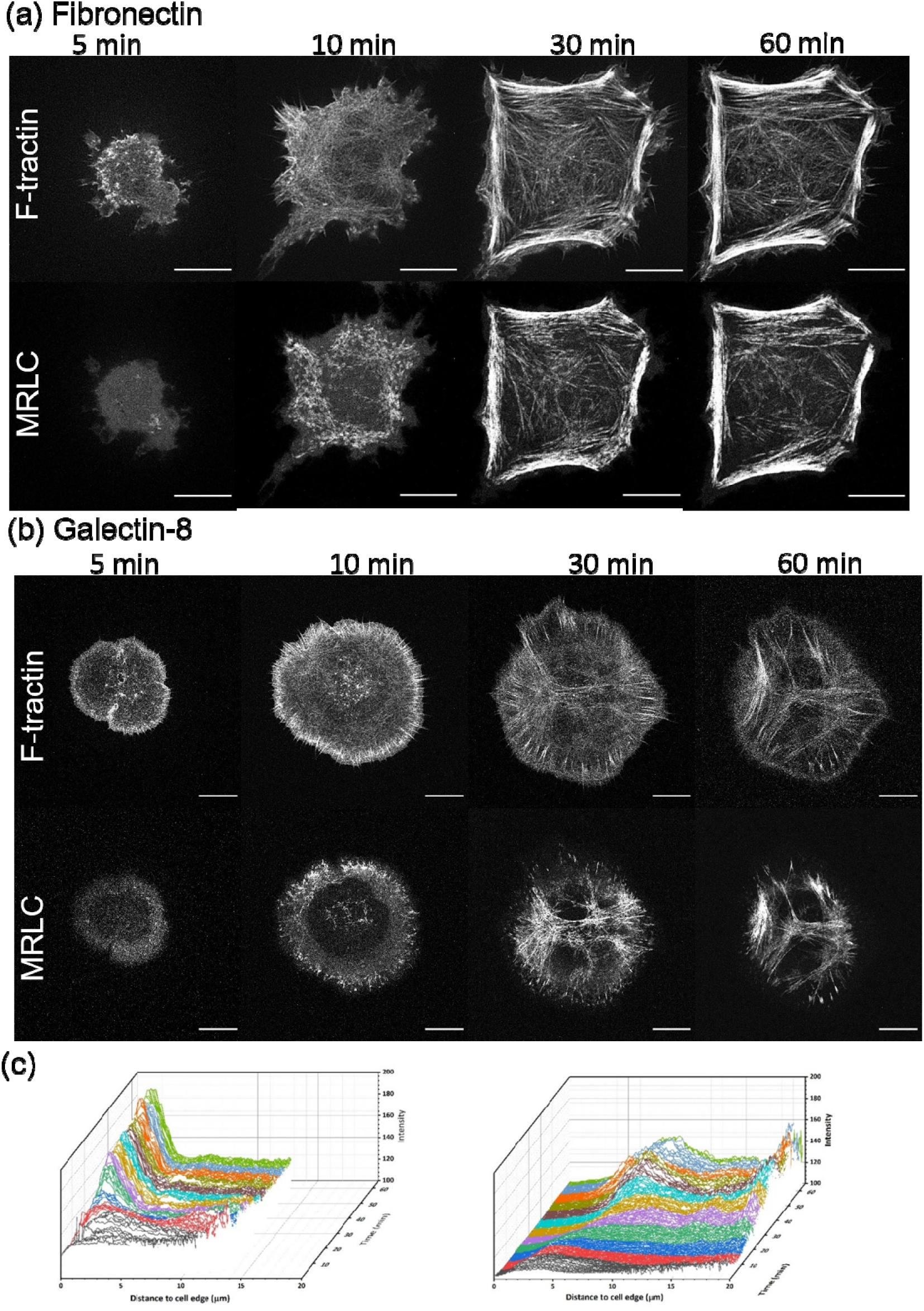
Actin and myosin II dynamics during cell spreading. Time course of spreading of cells transfected with tdTomato-F-tractin (actin) and GFP myosin II regulatory light chain (MRLC) on fibronectin (a) and galectin-8 substrates (b) (see also Movie 3 and Movie 4). Scale bar: 10 μm. Note formation of prominent actin and MRLC containing fibers on the periphery of cells spreading on fibronectin (a) and contracted actomyosin arrays of cells spreading on galectin-8 (b). Ten minutes after plating, numerous non-organized myosin filaments are seen on both substrates. (c) Total intensity of GFP-MRLC fluorescence as a function of the distance from the cell edge, in cells plated on fibronectin (left) and on galectin-8 (right). The intensity profiles corresponding to different time intervals are colored arbitrarily to facilitate visual assessment.

On galectin-8-coated substrates, cells displayed denser (though not necessarily longer) actin-positive filopodia than on substrates coated with fibronectin (Fig. 3b). The lamellipodia were enriched more strongly by F-actin than on fibronectin, and occupied the entire cell periphery in circular or petaloid cells. Tiny actin bundles were shown growing in a centripetal direction from the cell edges. At least some of these bundles appeared to constitute a continuation of filopodia. The myosin II filaments first appeared at the cell periphery, at a birth rate similar to those seen on the fibronectin-coated substrate (Movie 5). However, circumferential actomyosin bundles lying parallel to the cell edges were not detected in cells spreading on the galectin-8 substrates. In contrast to formation of the prominent peripheral actomyosin bundles, a star-like system of myosin II-enriched actin structures appeared in the central region of the cells (Fig. 3b; Movie 5). Quantification of the density of GFP-myosin II light chain containing filaments confirmed that on fibronectin-coated substrates, the myosin II filaments concentrate at the cell periphery, while on galectin-8-coated substrates, move toward the cell center (Fig. 3c).

### Differences in dynamics of focal adhesions in cells spreading on fibronectin and galectin-8

To investigate focal adhesion dynamics during the initial stages of cell spreading, we utilized HeLa cells stably transfected with YFP-paxillin, and also transfected with mCherry-Lifeact to visualize F-actin, and imaged the cells using TIRF microscopy. On fibronectin, the cells displayed the formation and maturation of focal adhesions (Fig. 4a; Movie 6), as described in many previous studies (Gardel et al., 2010; Geiger et al., 2009; Wolfenson et al., 2009). Maturation of focal adhesions is manifested by an increase in their area, as well as an increase in the fluorescence intensity (density) of major plaque proteins such as paxillin (Fig. 4a,c,e,f; Movie 6).

**Figure 4:**
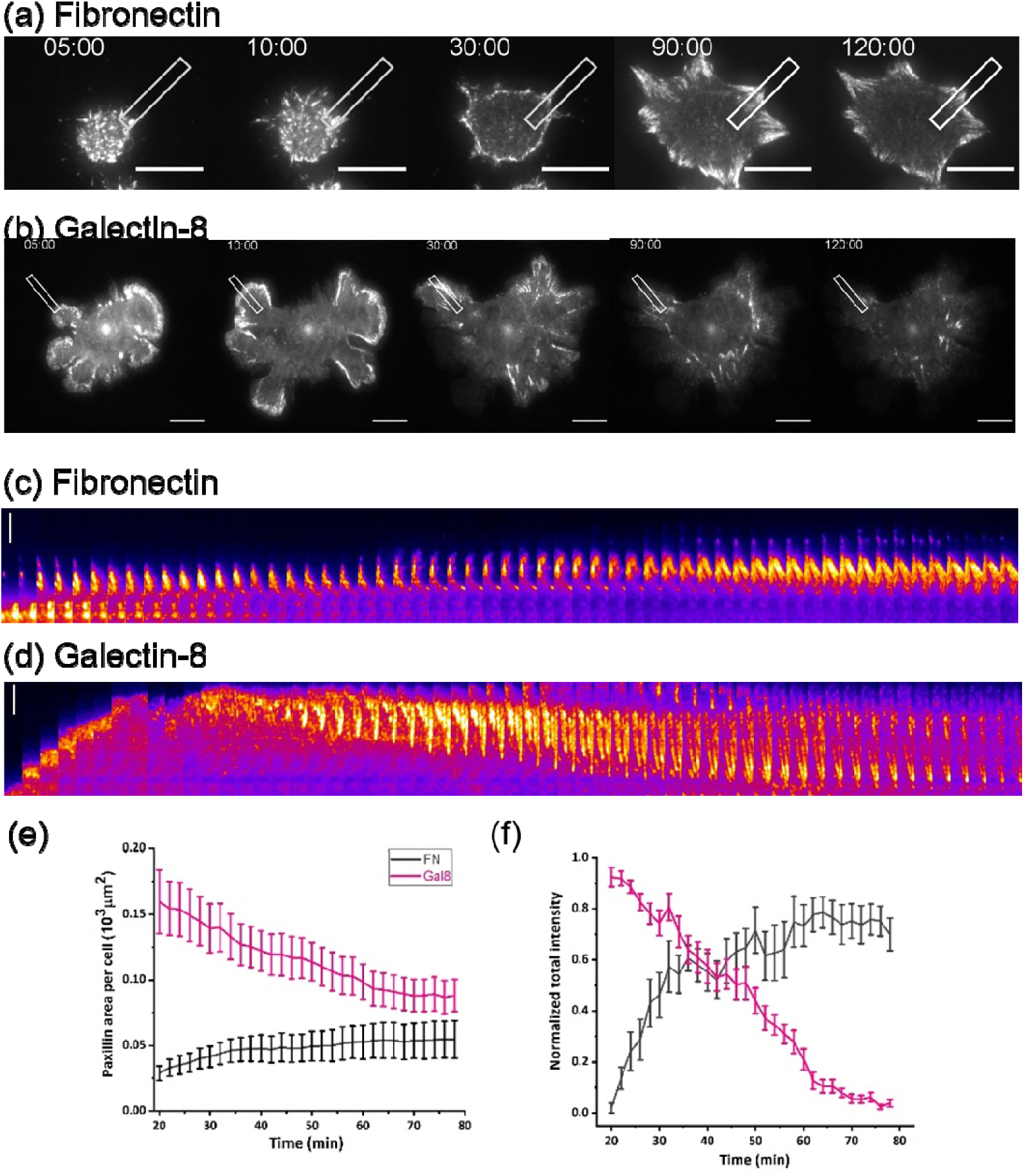
Paxillin dynamics during cell spreading on fibronectin and galectin-8. Sequences of images of cells stably expressing YFP-paxillin in the course of spreading on fibronectin (a) and galectin-8 (b) (see Movies 5 and 6). Scale bars: 15 μm. Note formation and growth of focal adhesions in (a) and formation, centripetal movement and disappearance of paxillin clusters in (b). (c)(d) High magnification kymographs showing paxillin dynamics in the rectangular strips (3 μm×l9 μm) drawn through the cells periphery perpendicular to the cell edge in cells plated on fibronectin (c) and on galectin-8 (d). Scale bars: 5 μm. Time intervals between each frame: 1 minute. (e)(f) Dynamics of the total area of paxillin clusters (e) and total intensity of YFP-paxillin fluorescence (f) per cell, for cells plated on fibronectin (black) and galectin-8 (pink). Error bar shows the standard error of mean (SEM). Number of assessed cells: N = 20.

On galectin-8, evolution of paxillin-enriched complexes proceeded in an entirely different manner. At the early stage of spreading, relatively large clusters of paxillin were located at the periphery of actin-rich lamellipodial protrusions (Fig. 4b; Movie 7; Supplementary Fig. 1). When the rate of lamellipodial extensions slowed, the paxillin clusters disintegrated into smaller patches, which moved centripetally and formed radially oriented chains associated with thin actin fibers (Fig. 4d; Supplementary Fig. 1). During the retrograde movement, paxillin clusters decreased in size and brightness, and gradually disappeared. Two hours after cell plating, most of the paxillin structures had vanished (Fig. 4b, d; Supplementary Fig. 1). Quantification of the areas and fluorescence intensity of paxillin clusters formed by the cells on galectin-8 as compared to fibronectin (Fig. 4e, f) clearly illustrate the differences in the dynamics of adhesion structures on these two substrates. Blocking the carbohydrate recognition domain (CRD) of galectin-8 by means of thiodigalactoside (TDG) (Kaufman and Lawless, 1980) did not change the projected cell area on fibronectin; neither did it inhibit formation of paxillin-positive focal adhesions (in fact, their size even increased slightly) (Supplementary Fig. 2a, c, d). However, treatment with TDG dramatically reduced the projected area of cells spreading on galectin-8, and diminished the size of paxillin clusters (Supplementary Fig. 2b, c, d). In agreement with Levy *et al*. (Levy et al., 2006), our experiments with galectin-8 deletion mutants revealed that both the N- and C-terminal carbohydrate-binding domains (CRD), as well as the extended hinge between these domains, are necessary to induce the full galectin-8 spreading phenotype characterized by extended lamellipodia formation. Cells only very weakly attached to galectin-8 lacking N-terminal CRD (Gal8-C) (Supplementary Fig. 2g). On galectin-8 lacking C-terminal CRD (Gal8-N) or the hinge (Gal8-Δhinge), the cells still attached, but mainly formed filopodia/retracting fibers, and not lamellipodia (Supplementary Fig. 2 h, i). In addition to the differences seen in the spreading and assembly of the actomyosin cytoskeleton and focal adhesions on fibronectin and galectin-8 coated substrates, we revealed that the strength of the adhesion that developed immediately after cells attached to the galectin-8 coated substrate was significantly higher than that seen after cells attached to the fibronectin substrate.

We assessed the forces required to detach cells from each substrate, using FluidFM technology in which the AFM cantilever was supplied with a microfluidic channel that permitted immobilization of the cell at the cantilever by applying negative pressure (Fig. 5a). The cell immobilized on the cantilever was allowed to contact the substrate, and the moment of initial contact was detected by cantilever deflection. Five minutes following the initial contact, the cells were detached from the substrate by uniaxial retraction of the cantilever. During this process, the deflection of the cantilever proportional to the applied force was recorded, and the maximum detachment force (MDF) was extracted as a representative parameter characterizing cell adhesion (Sancho et al., 2017). We found that the forces required to detach cells from the galectin-8 coated substrate were dramatically higher than the forces required to detach cells from the fibronectin-coated substrate (Fig. 5b). Furthermore, even five minutes after initial contact, the cell spreading area on galectin-8 was found to be somewhat larger than that seen on the fibronectin-coated substrate. The MDF per unit of cell spreading area was still much bigger for cells spreading on galectin-8 than for cells spreading on fibronectin (Fig. 5c).

**Figure 5:**
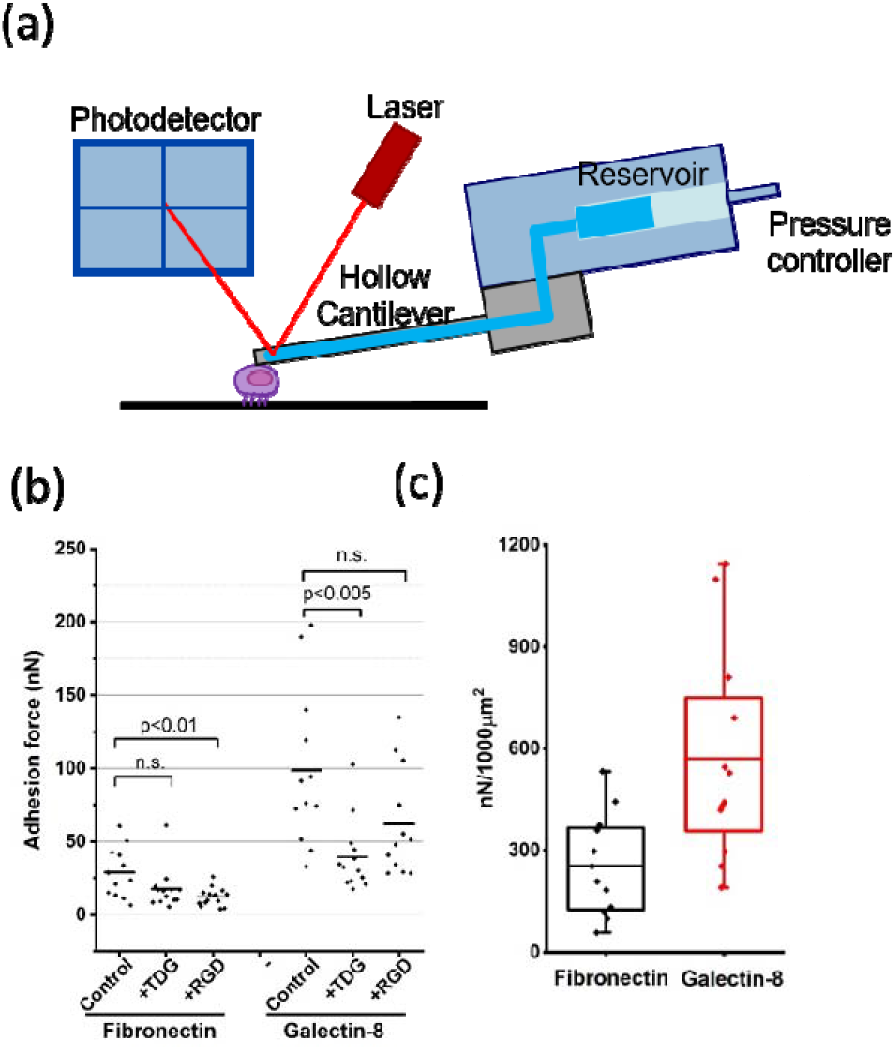
Cell-substrate adhesion forces on fibronectin and galectin-8 during early cell spreading. (a) A cartoon depicting the set-up of the microfluidic device used to measure adhesion forces. A pulling force is applied to the cell by a vacuum system integrated with the AFM cantilever. Adhesion force is defined as the minimal force sufficient to detach the cells from the substrate. (b) Measurement of cell-substrate adhesion forces in HelaJW cells five minutes following their plating on either fibronectin or galectin-8 in serum-free medium (control) or in the medium containing 20 mM TDG or 10 μg/ml RGD. The adhesion force was measured to be much stronger on the galectin-8 coated than on the fibronectin-coated substrate. Addition of TDG or RGD reduced the adhesion force on galectin-8- and fibronectin-coated substrates, respectively. (c) Adhesion forces on fibronectin and galectin-8 substrates, normalized per projected cell area (μm2). Note that the cell substrate adhesion force per unit of projected cell area on galectin-8 is significantly greater than that measured on fibronectin.

The addition of TDG reduced the force required for cell detachment from the galectin-8 coated, but not the fibronectin-coated substrate, while the addition of Arg-Gly-Asp (RGD) derivative with cysteine at its N-terminal (CGGGRGD) reduced the force required to detach cells from the fibronectin-coated substrate, but only weakly affected the force necessary to detach cells from the galectin-8 coated substrate (Fig. 5b).

### Effects of experimental manipulations with small Rho GTPases on cells spreading on galectin-8 and fibronectin-coated substrates

Small GTPases of Rho family are the master regulators of the actin cytoskeleton. Their activation downstream of interactions of cells with the extracellular matrix is thought to determine the processes of cell adhesion, spreading, and polarization upon the cell contact with the matrix. To elucidate the mechanism of cell response to fibronectin and galectin-8-coated substrates, we investigated the functions of three major Rho GTPases, RhoA, Rac1, and Cdc42, during cell spreading on these two substrates.

Half an hour following plating on fibronectin and galectin-8 coated substrates, RhoA and Rac1 activity was about 1.5-fold higher, and Cdc42 activity was about 2-fold higher, in cells on galectin-8 than those on fibronectin (Supplementary Fig. 3a, b). We assessed how activation or depletion of small Rho GTPases, as well as of their downstream targets, affect cell spreading on both substrates. The quantitative parameters chosen to characterize the cell spreading were: 1) cell projected area, reflecting bulk protrusion activity of lamellipodia; 2) the size of paxillin/vinculin-positive adhesion structures as a fraction of cell area; and 3) the number and average length of adherent cellular filopodia (Fig. 6).

**Figure 6:**
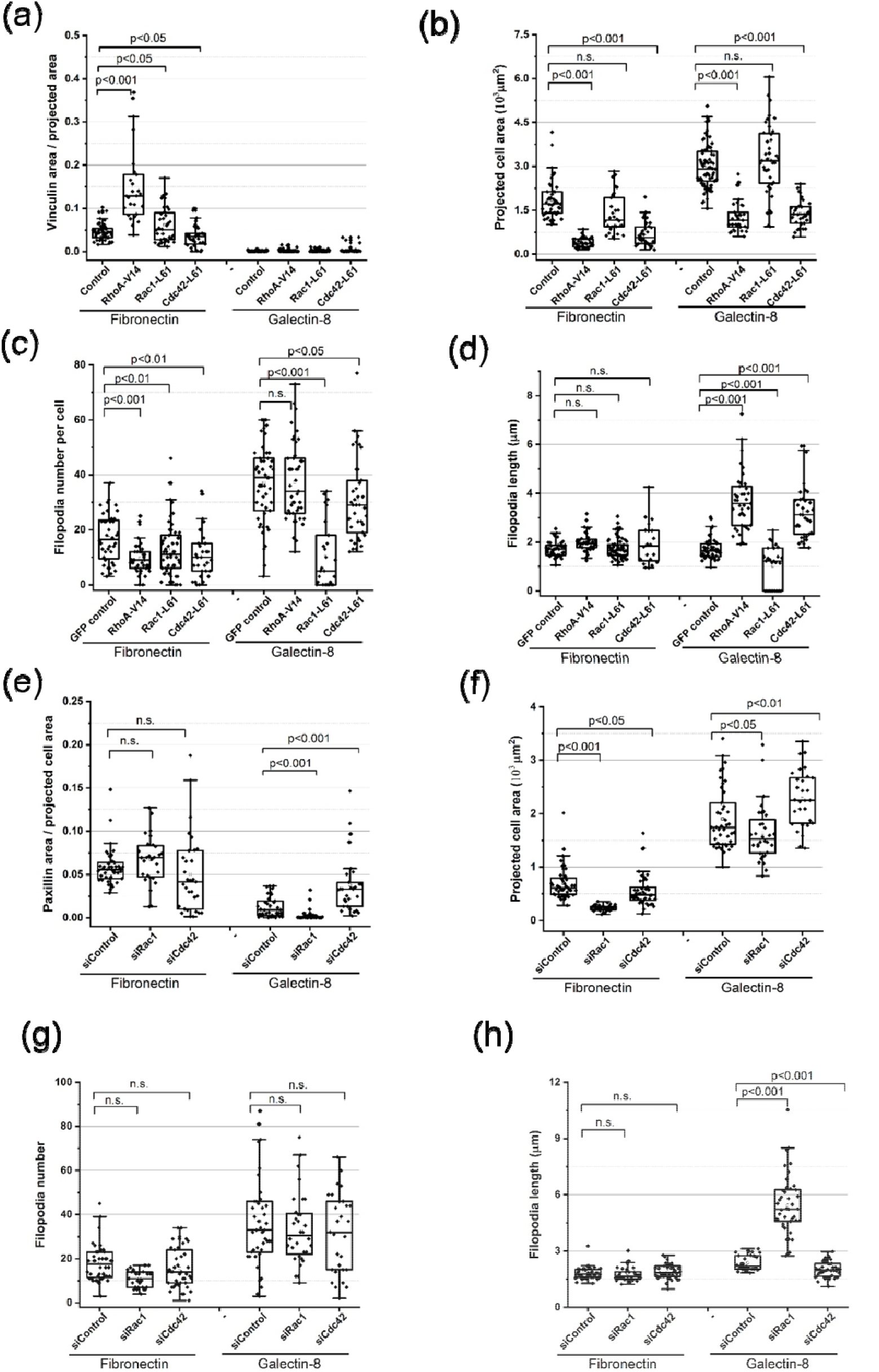
Effects of small Rho GTPases on cell spreading on fibronectin- and galectin-8 coated substrates. (a-d) Focal adhesions, projected cell area, and filopodia in control HelaJW cells, and in cells transfected with constitutively active small Rho GTPases, GFP-RhoA-V14, GFP-Rac1-L61, and GFP-Cdc42-L6l, assessed 2 hours after plating on fibronectin- and galectin-8 coated substrates, (e-h) Similar parameters assessed in control, Rac1 knock-down, and Cdc42 knock-down cells half an hour after plating. (a) and (e) Focal adhesion area as a fraction of total cell area; (b) and (f) Projected cell area; (c) and (g) Filopodia number per cell; (d) and (h) Average length of filopodia. Cells fixed and stained with TRITC-phalloidin and antibodies for vinculin (a-d) or paxillin (e-h) were used for morphometric measurements. Data are presented as box and whisker plots showing median values, upper and lower quartiles, maximum and minimum, and outliers (values which are 1.5 times larger than the upper or 1.5 times smaller than the lower quartiles). In all graphs, each dot corresponds to an individual cell.

Activation of RhoA by either expression of constitutively active RhoA-V14 (Fig. 6 a-d; Supplementary Fig. 3 c-j) or addition of the pharmacological activator CN03 (Cytoskeleton, Inc.) (Flatau et al., 1997; Schmidt et al., 1997) (Supplementary Fig. 4) significantly promoted formation of vinculin- or paxillin-positive focal adhesions and linear actin stress fibers on the fibronectin-coated substrate, but did not protect vinculin/paxillin clusters from disappearance on the substrates coated with galectin-8 (Fig. 6a; Supplementary Fig. 3c-j; Fig. 4a-h, j). The projected area of cells on fibronectin decreased upon RhoA activation. On galectin-8, despite RhoA activation, the projected area always remained larger than that on fibronectin (Fig. 6b; Supplementary Fig. 4i). Constitutively active RhoA promoted formation of filopodia on the galectin-8 coated substrate (Fig. 6c, d; Supplementary 3j; Movie 8). These stimulated filopodia were much longer than those seen in control cells (Fig. 6d), and often displayed triangular actin-enriched “pedestals” at their bases (Supplementary Fig. 3j’).

Expression of constitutively active Rac1-L61 resulted in a circular cell shape, promoting lamellipodia formation on both substrates (Supplementary Fig. 3k-n; Movie 9). On the fibronectin-coated substrate, it led to formation of radial and circumferential actin bundles associated with enlarged focal adhesions (Fig. 6a; Supplementary Fig. 3k, l), while on the galectin-8 coated substrate, formation of focal adhesions remained completely suppressed, and actin was enriched in lamellipodia (Fig. 6a; Supplementary Fig. 3m, n). The difference between the spreading area of cells on fibronectin and galectin-8 coated substrates did not decrease upon Rac1 activation (Fig. 6b). At the same time, constitutively active Rac1 strongly suppressed filopodia formation on galectin-8 (Fig. 6c, d; Supplementary Fig. 3n, n’).

Expression of constitutively active Cdc42-L61 decreased lamellipodia formation and spreading area on both substrates (Fig. 6b; Supplementary Fig. 3o-r). It did not rescue the formation of focal adhesions and stress fibers on the galectin-8 coated substrate (Fig. 6a; Supplementary Fig. 3q, r). Active Cdc42-L61 did not appear to increase filopodia number or length on the fibronectin-coated substrate, but augmented filopodia length on the galectin-8 coated substrate (Fig. 6c, d; Supplementary Fig. 3r, r’). These stimulated filopodia displayed actin-enriched “pedestals,” similarly to filopodia stimulated by RhoA (Supplementary 3j, j’).

We then studied the effects of Rac1 and Cdc42 knock-down at the early stages of cell spreading (30 minutes after seeding) on fibronectin and galectin-8 coated substrates. On fibronectin, knock-down of either Rac1 or Cdc42 did not significantly affect the fraction of cell area occupied by paxillin-positive adhesions (Fig. 6e). Paxillin clusters formed at the initial stages of spreading on the galectin-8 coated substrate were completely abolished in Rac1-knock-down cells, while in Cdc42 knock-down cells, these clusters became larger than in control cells on galectin-8 (Fig. 6e; Supplementary Fig. 5). Knock-down of Rac1 considerably decreased the cell spreading area on fibronectin (about two folds), but only slightly reduced the spreading on galectin-8 (Fig. 6f; Supplementary Fig. 5e, g), while knock-down of Cdc42 slightly decreased the cell spreading area on fibronectin, but augmented it on galectin-8 (Fig. 6f; Supplementary Fig. 5i, k). Clear differences were noted between the projected cell area on the fibronectin and galectin-8 coated substrates for both control cells, and Rac1 and Cdc42 knock-down cells (Fig. 6f). Rac1 knock-down did not significantly change filopodia number and length on fibronectin, but strongly increased filopodia length on the galectin-8 coated substrate (Fig. 6g, h; Supplementary Fig. 5g), while knock-down of Cdc42 did not affect filopodia number, and only slightly reduced filopodia length on galectin-8 coated substrates (Fig. 6g, h; Supplementary Fig. 5k).

Among downstream effectors of small GTPases, we investigated the effects of activation or depletion of actin polymerization regulators, the Arp2/3 complex and the formins mDia2 and FMNL2, on early cell spreading on galectin-8. In addition, we assessed the effects of pharmacological inhibition of the RhoA target, Rho kinase (ROCK). Expression of constitutively active constructs of mDia2 and FMNL2 decreased projected cell area (Fig. 7a; Supplementary Fig. 6a-c), did not increase filopodia number, but increased filopodia length (Fig. 7b, c). Consistently, knock-down of mDia2 and FMNL2 increased projected cell area, and suppressed formation of filopodia (Fig. 7d, e). These effects contrasted with those caused by depletion of the Arp2/3 complex by Arp2 knock-down, the latter of which significantly decreased the projected cell area, presumably by inhibition of lamellipodia (Fig. 7d) and enhanced formation of filopodia (Fig. 7e, f). Finally, inhibition of ROCK by specific inhibitor Y27632 increased the projected cell area on both fibronectin and galectin-8. However, the projected cell area on galectin-8 remained much larger than that seen on fibronectin (Supplementary Fig. 7a). In addition, Y27632 treatment reduced the area of paxillin clusters on both substrates (Supplementary Fig. 7b), and the number of filopodia on galectin-8 (Supplementary Fig. 7c).

**Figure 7:**
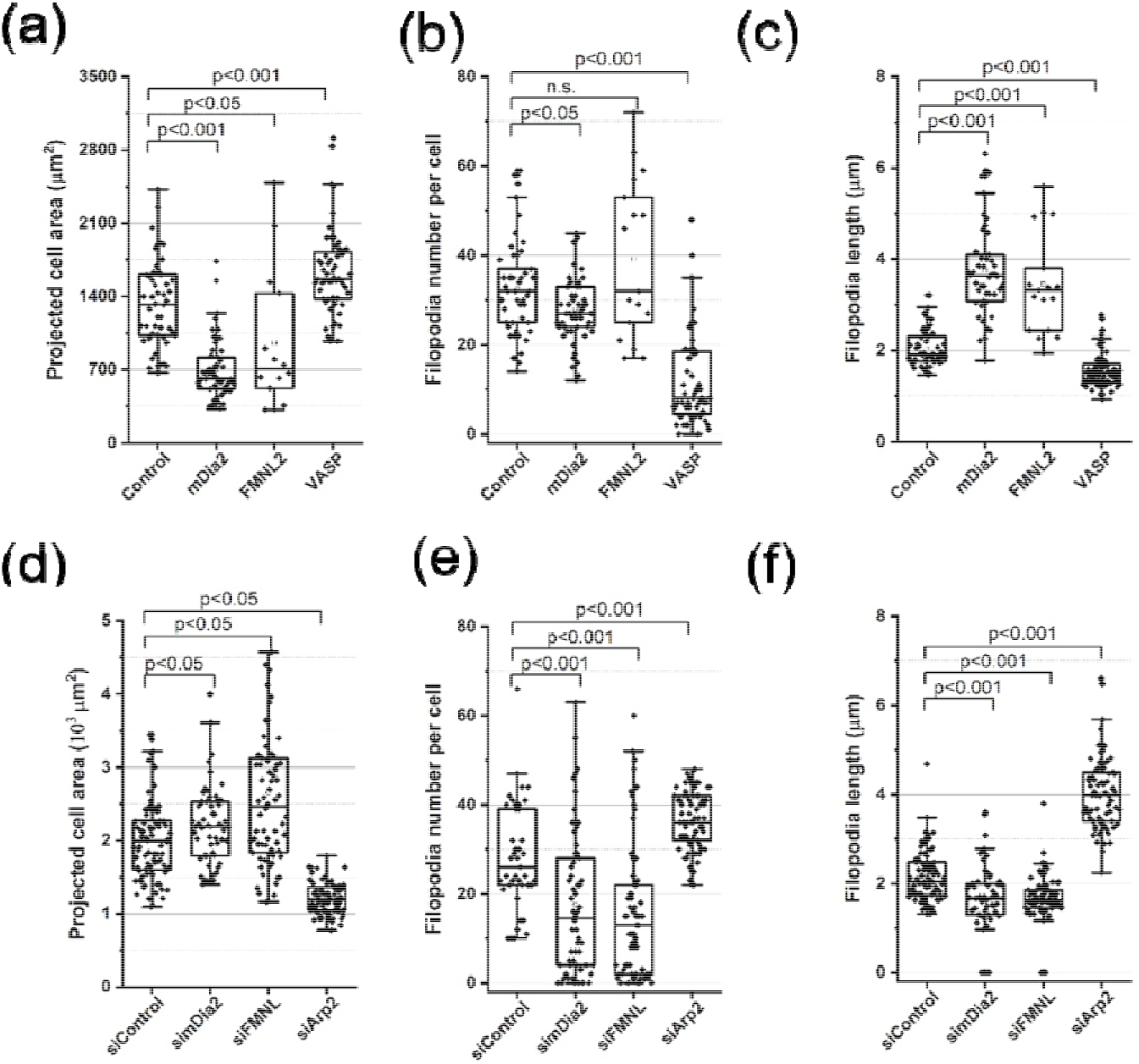
Effects of actin polymerization regulators on cell spreading on a galectin-8 coated substrate. (a-c) Control cells and cells overexpressing GFP-mDia2, GFP-FMNL2, and GFP-VASP. (d-f) Control cells and cells expressing siRNAs against mDia2, FMNL2, and Arp2. Cells were plated on galectin-8 coated substrates in serum-free medium, and fixed twenty minutes following plating. Cells were stained with TRITC-phalloidin. Morphometric measurements and presentation of results were performed as described in the legend to Figure 6.

### Combined effects of fibronectin and galectin-8 on cell spreading, focal adhesion and filopodia formation

*In vivo,* cells only rarely, if at all, encounter an extracellular matrix consisting of one type of protein. As a rule, cell behavior is determined by a complex mixture of extracellular ligands. Here, we studied a simplified situation, analyzing cellular response to a mixture of two types of matrix proteins, fibronectin and galectin-8, employed at different ratios. A time point of four hours was chosen to assess the cells at maximal spreading. At this time point, the average spreading area on the fibronectin-coated substrate approached a maximal value of ~1700 μm^2^. The substrate coated with a mixture of fibronectin at a maximal concentration (25 μg/ml), and an increasing concentration of galectin-8 resulted in a gradual increase in cell spreading area up to ~2800 μm^2^, seen at the maximal concentration of galectin-8 (25 μg/ml). Remarkably, a gradual decrease in the concentration of fibronectin mixed with the maximal concentration of galectin-8 led to an additional small, though significant increase of the projected cell area (Fig. 8a, b). Thus, galectin-8 strongly regulates cell spreading in a positive manner, even when mixed with the maximal concentration of fibronectin; fibronectin, on the other hand, exerts a slight negative effect on cell spreading at the maximal concentration of galectin-8.

**Figure 8:**
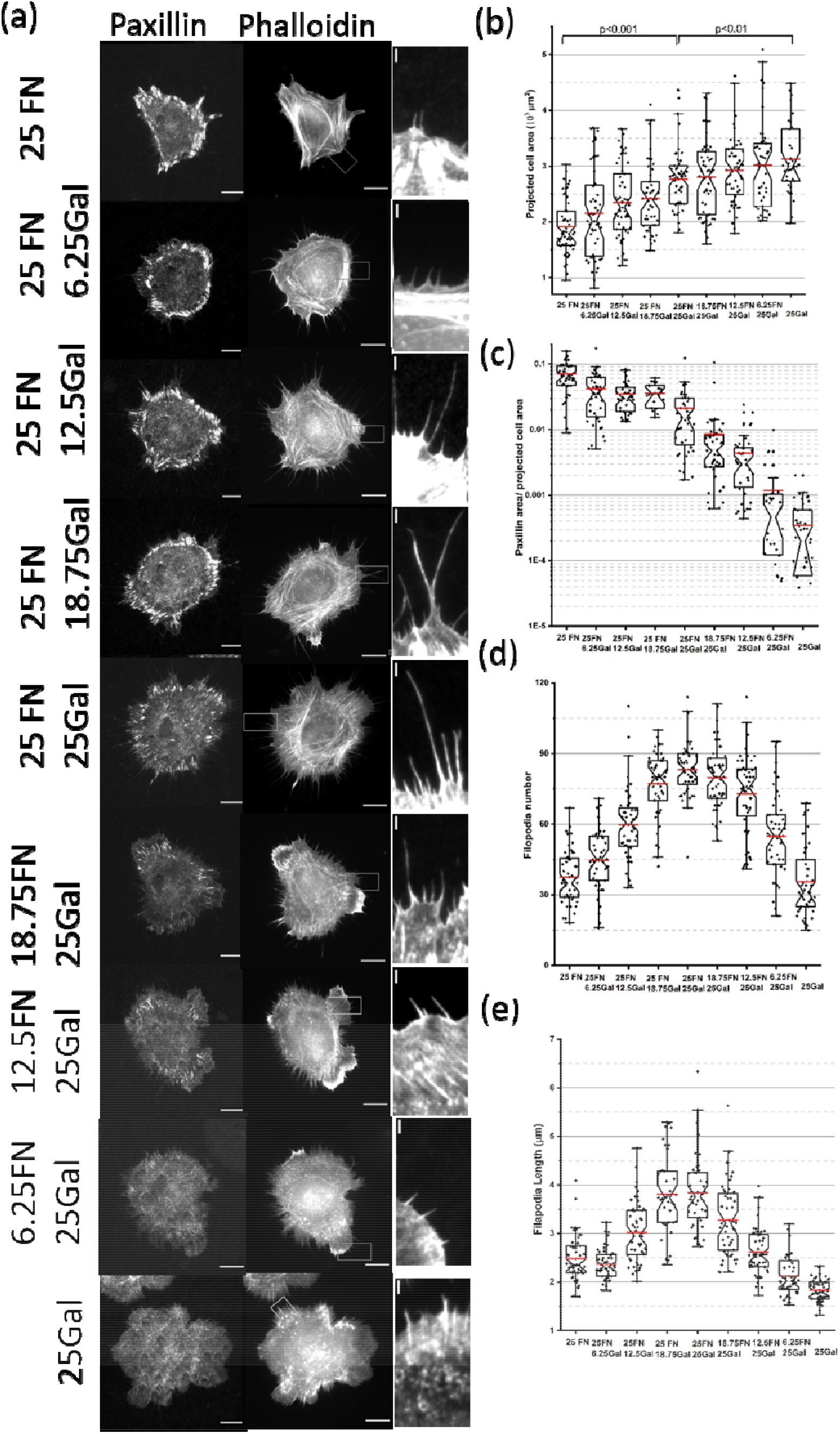
Cell spreading on composite matrices comprising different fibronectin/galectin-8 ratios. (a) Focal adhesions visualized by YFP-paxiIlin (left column), and F-actin visualized by TRITC-phaIloidin staining (centerl and right columns) in cells fixed 4 hours following plating on substrates coated with different combinations of fibronectin (FN) and galectin-8 (Gal), as indicated. The right column displays the boxed areas seen in the images shown in the center column, at high magnification. (b-e) Quantification of spreading area (b), total paxillin adhesion area (c), filopodia number (d) and average length per cell (e). Each dot corresponds to an individual cell; N ≥40 cells were assessed under each experimental condition. Results are presented as a box and whisker plot, as in Figures 6 and 7.

The size of paxillin-containing complexes also changed in response to different ratios between fibronectin and galectin-8. Focal adhesions developed to the maximum extent at the highest concentration of fibronectin, and only slightly decreased (if at all) when galectin-8 concentrations were increased (Fig. 8a, c). A decrease in fibronectin concentration at the maximal concentration of galectin-8, produced a decrease in the area of focal adhesions, which, however, still existed. The focal adhesions on pure galectin-8 substrate disappeared at this time point (four hours after seeding) (Fig. 8a, c). Thus, galectin-8 does not support the formation of focal adhesions, but also does not inhibit formation of focal adhesions in the presence of fibronectin. Even small concentrations of fibronectin can promote the formation of some focal adhesions in the presence of the maximal concentration of galectin-8.

Unlike the projected cell area, which responded mainly to changes in galectin-8 concentration, and focal adhesions, which responded mainly to changes in fibronectin concentration, filopodia formation was sensitive to the concentrations of both ligands. On the substrate consisting solely of fibronectin, the number of filopodia per cell was relatively low (37 per cell, on average). Addition of increasing concentrations of galectin-8 resulted in a marked increase in filopodia number (above 80 per cell) after four hours of spreading (Fig. 8a, d). The gradual decrease in fibronectin concentration, under conditions of a constant, maximal galectin-8 concentration, also led to a dramatic decrease in filopodia number (Fig. 8a, d). Altogether, the dependence of filopodia number on the concentrations of these two ligands demonstrates their synergistic effect, as the maximal number of filopodia is approached when maximal concentrations of fibronectin and galectin-8 are combined, while a decrease of either fibronectin or galectin-8 strongly decreased filopodia number. Of note, at optimal concentrations of fibronectin and galectin-8, thus maximizing the filopodia number, the mean filopodia length was also significantly higher than at non-optimal concentrations of these ligands (Fig. 8e).

## Discussion

Here, we demonstrated that the cells spreading on substrates covered by fibronectin and galectin-8 are strikingly different. The elementary processes comprising the spreading include formation of cell extensions, lamellipodia and filopodia, and adhesions of these extensions to the substrate. These processes are brought about by reorganizations of the actomyosin cytoskeleton. Lamellipodial and filopodial extensions are filled with an Arp2/3 nucleated branched actin network and formin nucleated actin bundles, respectively (Blanchoin et al., 2014; Campellone and Welch, 2010; Chhabra and Higgs, 2007; Le Clainche and Carlier, 2008). Focal adhesion plaques are built of actin filaments and associate with the actin bundles, which transmit actomyosin-generated forces (Medalia and Geiger, 2010; Xia and Kanchanawong, 2017). The spreading that results is seen in an increase of projected cell area, and in the development of a characteristic cell shape and adhesion pattern. The processes of cell spreading on a fibronectin-coated rigid planar substrate have been well studied. The spreading area increases due to formation of lamellipodia, which trigger formation and maturation of focal adhesions associated with the system of actomyosin bundles (stress fibers) (Small et al., 1998; Wolfenson et al., 2014). Subsequent reorganization of the stress fibers in turn determines the elongation and polarization of initially radial symmetric cells (Prager-Khoutorsky et al., 2011). On the substrate coated with galectin-8, all stages of this process differ. Formation of lamellipodia proceeds more persistently, without pause and retraction episodes typically seen in cell spreading on fibronectin. Several lamellipodia are usually formed simultaneously at 3-8 segments of the cell periphery, resulting in development of a characteristic “petaloid” cell contour. Remarkably, physical adhesion of cells to galectin-8 is much stronger than to the fibronectin substrate, as demonstrated in our measurement of forces per unit of cell spreading area required to detach cells from the substrates. Five minutes after initial contact of the cell with the substrate, such forces were 2-fold higher on the galectin-8-than on the fibronectin-coated substrate.

Rapidly extending and strongly adherent lamellipodia on galectin-8-coated substrates were, however, unable to form focal adhesions. While on the fibronectin substrate, cells formed numerous dot-like nascent adhesions which eventually matured into the typical elongated focal adhesions associated with stress fibers (Burridge, 2017; Geiger et al., 2009), on galectin-8, cells initially formed large paxillin- and vinculin-enriched clusters at the cell periphery, which then moved centripetally and eventually disappeared. Accordingly, organization of the actomyosin fiber system proceeded differently on each of these two substrates. The peripheral actomyosin bundles, which exert traction forces on focal adhesions typical of cells spreading on fibronectin (Burridge and Guilluy, 2016; Kassianidou and Kumar, 2015; Tojkander et al., 2012), never formed on galectin-8. Myosin II filaments still intensely assemble at the cell periphery during cell spreading, but instead of incorporating into circumferential actin bundles, they form several star-like contraction foci randomly located in the central part of the cell. Finally, cell spreading on galectin-8 involved the formation of numerous filopodia extending from the edges of lamellipodia. While cells spreading on fibronectin could also form filopodia, especially at the early stages of attachment, such filopodia often failed to adhere to the substrate, and rapidly retracted. Formation of numerous adherent filopodia at advanced stages of spreading was typically seen in cells spreading on galectin-8.

A potentially important mechanism underlying the differences in cells spreading seen on fibronectin and galectin-8 coated substrates could depend on varying downstream signals from the adhesion receptors. Since the receptors involved in cell adhesion to galectin-8 are numerous (Bieniasz-Krzywiec et al., 2019; Cárcamo et al., 2006; Cueni and Detmar, 2009; Elola et al., 2014; Fernández et al., 2016; Ferragut et al., 2019; Hadari et al., 2000; Troncoso et al., 2014; Vinik et al., 2015) and only partially identified, it is difficult to analyze the differences between these signaling pathways in full detail. Therefore, we chose to focus on the downstream level of signaling; namely, activation of the small G-proteins of the Rho family, and their major cytoskeletal targets.

It is well-established that on fibronectin-coated substrates, spreading is initiated by activation of Rac1 and Cdc42, which in turn triggers Arp2/3-dependent branched actin polymerization (Devreotes and Horwitz, 2015; Price et al., 1998). The integrin-mediated activation of RhoA at the later stages of cell spreading on rigid fibronectin-coated substrates promotes myosin-II driven actomyosin contractility, which restricts further spreading and triggers maturation of focal adhesions (Burridge and Guilluy, 2016; Burridge et al., 2019; Ren et al., 1999). Attachment of cells to a galectin-8 coated substrate apparently also triggers activation of the small Rho GTPases (Cárcamo et al., 2006; Diskin et al., 2012). Our pull-down experiments reveal that Rac1 and RhoA activities in cells spreading on galectin-8 were slightly increased, compared to cells spreading on fibronectin. This difference was even more pronounced for Cdc42 activity. While a quantitative estimation of the ratios between the activities of the small GTPases in the course of spreading using the pull-down technique is problematic, we hypothesize that phenotypic differences between cells on fibronectin and galectin-8 coated substrates are due at least in part to changes in the balance between the activities of major Rho GTPases.

To elucidate the effects of these GTPases on phenotypical features of spreading cells, we performed several experiments involving activation and depletion of these GTPases. On galectin-8, the area of cell spread can be reduced by overexpression of constitutively active RhoA or Cdc42, while knock-down of Cdc42 slightly increases overall cell spreading. These effects could be explained by suppression of lamellipodia formation and/or increase of overall cell contractility by RhoA and Cdc42. Indeed, both RhoA and Cdc42 can activate myosin II light chain phosphorylation via their immediate targets ROCK and MRCK, respectively (Wilkinson et al., 2005; Zhao and Manser, 2015). Accordingly, we showed that inhibition of ROCK also increased the cell projected area on galectin-8.

Notably, manipulations affecting activities of RhoA/ROCK and Cdc42, which reduce/augment cell projected area, exert the opposite effects on filopodia formation. Indeed, constitutively active Cdc42 and RhoA increase filopodia number and/or length on galectin-8, while knockdown of Cdc42 or inhibition of ROCK decrease it. At the same time, Rac1 is antagonistic to filopodia formation on galectin-8, since its constitutively active mutant strongly reduces filopodia number and length, while its depletion augments filopodia. Effects of altered activity of small G-proteins on cell projected area and filopodia formation are consistent with the effects of their immediate targets, regulators of actin polymerization, the Arp2/3 complex and formins. Overexpression of two major formins, mDia2 and FMNL2, which are downstream targets of RhoA and Cdc42 (Kühn and Geyer, 2014), activated filopodia formation but decreased cell projected area. Knock-downs of mDia2 and FMNL2 decreased filopodia number and increased cell projected area. Moreover, knock-down of Arp2, leading to suppression of Arp2/3 actin polymerization activated by Rac1, decreased the projected cell area, but promoted formation of filopodia, augmenting their number and length, in agreement with previous studies (Innocenti, 2018; Steffen et al., 2014; Swaney and Li, 2016).

Thus, the overall pattern of formation of actin-rich extensions and cell spreading on galectin-8 coated substrates likely arises out of the interplay between two competing processes: Arp2/3-driven assembly of the branched actin network, resulting in formation and extension of lamellipodia, and formin-driven assembly of filopodia actin cores. These two processes are antagonistic, since they compete for the same pool of monomeric actin. Our data would suggest that while Arp2/3-driven actin polymerization in lamellipodia is mainly activated by Rac1, consistent with numerous data obtained in other systems (Innocenti, 2018; Steffen et al., 2014), formin-driven actin polymerization in filopodia is activated not only by Cdc42, which is broadly accepted, but also by the RhoA-ROCK signaling axis, perhaps via the activation of mDia2 (Staus et al., 2011). RhoA can also extend the lifespan of filopodia through myosin IIA-dependent promotion of filopodia adhesion to the extracellular matrix (Alieva et al., 2019). It is interesting that the competition between Rac1-driven lamellipodia and Cdc42/RhoA-driven filopodia seems to be more pronounced on the galectin-8 coated, rather than on the fibronectin-coated substrate. Indeed, on galectin-8, activation of Rac1 leads to the almost complete disappearance of filopodia, while down-regulation of Rac1 results in formation of numerous long filopodia. These effects are not seen on fibronectin.

Importantly, the differences between cell spreading area on galectin-8 and on fibronectin cannot be fully explained by changes in activity of small Rho GTPases and their targets. Even though the projected area of cells on galectin-8 can be reduced upon activation of RhoA, Cdc42, and formins as well as upon inhibition of Rac1 and the Arp2/3 complex, it still always exceeded the projected area on fibronectin. This suggests that other factors, most probably the strength of physical adhesions detected in our study, should be also taken into consideration.

Another difference between the phenotypes of cells spreading on galectin-8 and fibronectin substrates lies in the inability of cells on galectin-8 to assemble into mature focal adhesions and associated stress fibers. Instead, these cells already form large paxillin- and vinculin-containing clusters at the early stages of spreading (less than twenty minutes). These clusters are associated with the actin cytoskeleton, since they are moving centripetally, apparently pushed by the radial actin bundles in the lamellipodial “petals” of cells on galectin-8. This association did not, however, lead to maturation of the paxillin clusters into stable adhesion structures; rather, they disappeared 2-4 hours after cell plating.

The elimination of paxillin clusters in cells on galectin-8 cannot be prevented by constitutively active RhoA, Cdc42 or Rac1. At the same time, galectin-8 does not strongly inhibit the formation of focal adhesions if fibronectin is present on the substrate. The highest concentration of galectin-8 only slightly reduced the area of focal adhesions in the presence of fibronectin. Accordingly, even the smallest concentration of fibronectin could induce the formation of some focal adhesions at the highest concentration of galectin-8. Provided that galectin-8 interacts with several integrins (Cárcamo et al., 2006; Diskin et al., 2009; Elola et al., 2014; Hadari et al., 2000), it is not immediately clear why contact with galectin-8, coupled with activation of RhoA, are not sufficient to elicit formation of mature focal adhesions, in and of themselves. One possible explanation could be that the interaction of galectin-8 with glycosylated integrins differs sterically from interactions of fibronectin with integrins, and cannot promote proper integrin activation and the subsequent assembly of focal adhesions. It might also be that adhesion to galectin-8 triggers the formation of a special type of molecular cluster qualitatively different from classical focal adhesions, as well as other types of integrin-mediated adhesions.

Cell spreading on mixtures of fibronectin and galectin-8 in varying proportions revealed the ability of cells to combine signals produced by contacts with different types of the matrix. Galectin-8 increased the projected area in a concentration-dependent fashion, in the presence of the highest concentration of fibronectin. The effect of the mixture of matrix proteins on cell spreading area is dominated by galectin-8, and only slightly negatively regulated by fibronectin. Similarly, the effect of the matrix protein mixture on focal adhesion formation is dominated by fibronectin, which induces these structures in a concentration-dependent manner, in the presence of the highest concentration of galectin-8. The highest concentration of galectin-8 only slightly reduced the area of focal adhesions in the presence of the highest concentration of fibronectin. Unlike spreading area and focal adhesions, formation of filopodia on the mixed fibronectin-galectin-8 substrate markedly exceeded that seen on either individual substrate. Both filopodia number and length increase upon addition of even a small amount of one ligand to the maximal concentration of another ligand. Maximal filopodia formation was observed at the maximal concentrations of both ligands. This finding demonstrates the possible synergistic effects of different components of the extracellular matrix, which may play important roles in cell adhesion and migration *in vivo*.

In particular, formation of filopodia was shown to correlate with cancer cell metastasis (Arjonen et al., 2011; Jacquemet et al., 2015). This could explain why excessive production of galectin-8 could augment the metastatic capacity of cancer cells (Gentilini et al., 2017; Reticker-Flynn et al., 2012; Shatz-Azoulay et al., 2020).In summary, we demonstrated in this study that a galectin-8 coated substrate induces a cell adhesion response that differs from that induced by fibronectin. Cells on galectin-8 spread more rapidly and persistently, and approach a larger projected area than seen on fibronectin. This results from the increased efficiency of lamellipodia extension, which in turn depends on Arp2/3 activation as well as on stronger physical adhesions between cell receptors and the galectin-8 ligand, but is negatively regulated by RhoA, Cdc42 and formins. Cells plated on a galactin-8 coated substrate cannot, however, form persistent mature focal adhesions and an associated system of actin stress fibers. RhoA-induced myosin II filaments in cells spreading on the galectin-8 substrate do not assemble into transverse arcs and ventral stress fibers typical of cells on fibronectin. Rather, a characteristic feature of spreading on galectin-8 entails the robust formin-dependent formation of adherent filopodia triggered by Cdc42 and RhoA, and strongly opposed by Rac1-Arp2/3. This galectin-8 induced filopodia formation is synergistically activated by fibronectin, so that filopodia number and length on the substrate coated with a mixture of the ligands dramatically exceed those on each type of ligand separately. Such synergistic effects may play an important role in the cellular response to composite matrices *in vivo*.

## Acknowledgement

We thank Dr. Ed Manser (A-star, Singapore) for the FMNL2 formin construct.

B.G. is grateful to the Israel Science Foundation (personal grants and precision medicine grants) and to the Minerva Center at the Weizmann Institute “Aging, from Physical Materials to Human Tissues” for their support. A.D.B. acknowledge the support from the Singapore Ministry of Education Academic Research Fund Tier 2 (MOE Grant No: MOE2018-T2-2-138), the National Research Foundation, Prime Minister’s Office, Singapore, and the Ministry of Education under the Research Centres of Excellence programme through the Mechanobiology Institute, Singapore (ref no. R-714-006-006-271), and Singapore Ministry of Education Academic Research Fund Tier 3 MOE grant no. MOE2016-T3-1-002. A.D.B. and B.G. also acknowledge the support from a Maimonides Israeli–France grant (Israeli Ministry of Science Technology and Space), and EU Marie Skłodowska-Curie Network InCeM (project ID 642866) at the Weizmann Institute of Science. W.L. acknowledge the support from from the European Union’s Horizon 2020 research and innovation programme under the Marie Sklovowska-Curie grant agreement no. 642866. A.S. and J.G. acknowledge financial support from the ERC (European Research Council) under Grant Number 617989

## References

Adams, J.C., N. Kureishy, and A.L. Taylor. 2001. A role for syndecan-1 in coupling fascin spike formation by thrombospondin-1. J Cell Biol. 152:1169–1182.

Adams, J.C., and M.A. Schwartz. 2000. Stimulation of fascin spikes by thrombospondin-1 is mediated by the GTPases Rac and Cdc42. J Cell Biol. 150:807–822.

Alieva, N.O., A.K. Efremov, S. Hu, D. Oh, Z. Chen, M. Natarajan, H.T. Ong, A. Jégou, G. Romet-Lemonne, J.T. Groves, M.P. Sheetz, J. Yan, and A.D. Bershadsky. 2019. Myosin IIA and formin dependent mechanosensitivity of filopodia adhesion. Nat Commun. 10:3593–3593.

Alonso, F., P. Spuul, T. Daubon, I. Kramer, and E. Génot. 2019. Variations on the theme of podosomes: A matter of context. Biochimica et biophysica acta. Molecular cell research. 1866:545–553.

Antonio, C., S. Jamie, and K. Ender. 2012. Decision Forests: A Unified Framework for Classification, Regression, Density Estimation, Manifold Learning and Semi-Supervised Learning. now. 1 pp.

Arjonen, A., R. Kaukonen, and J. Ivaska. 2011. Filopodia and adhesion in cancer cell motility. Cell Adh Migr. 5:421–430.

Bai, M., B. Harfe, and P. Freimuth. 1993. Mutations that alter an Arg-Gly-Asp (RGD) sequence in the adenovirus type 2 penton base protein abolish its cell-rounding activity and delay virus reproduction in flat cells. J Virol. 67:5198–5205.

Berg, S., D. Kutra, T. Kroeger, C.N. Straehle, B.X. Kausler, C. Haubold, M. Schiegg, J. Ales, T. Beier, M. Rudy, K. Eren, J.I. Cervantes, B. Xu, F. Beuttenmueller, A. Wolny, C. Zhang, U. Koethe, F.A. Hamprecht, and A. Kreshuk. 2019. ilastik: interactive machine learning for (bio)image analysis. Nature methods. 16:1226–1232.

Bieniasz-Krzywiec, P., R. Martín-Pérez, M. Ehling, M. García-Caballero, S. Pinioti, S. Pretto, R. Kroes, C. Aldeni, M. Di Matteo, H. Prenen, M.V. Tribulatti, O. Campetella, A. Smeets, A. Noel, G. Floris, J.A. Van Ginderachter, and M. Mazzone. 2019. Podoplanin-Expressing Macrophages Promote Lymphangiogenesis and Lymphoinvasion in Breast Cancer. Cell metabolism. 30:917–936.e910.

Blanchoin, L., R. Boujemaa-Paterski, C. Sykes, and J. Plastino. 2014. Actin dynamics, architecture, and mechanics in cell motility. Physiological reviews. 94:235–263.

Bonnans, C., J. Chou, and Z. Werb. 2014. Remodelling the extracellular matrix in development and disease. Nat Rev Mol Cell Biol. 15:786–801.

Burridge, K. 2017. Focal adhesions: a personal perspective on a half century of progress. The FEBS journal. 284:3355–3361.

Burridge, K., and C. Guilluy. 2016. Focal adhesions, stress fibers and mechanical tension. Exp Cell Res. 343:14–20.

Burridge, K., E. Monaghan-Benson, and D.M. Graham. 2019. Mechanotransduction: from the cell surface to the nucleus via RhoA. Philos Trans R Soc Lond B Biol Sci. 374:20180229–20180229.

Campellone, K.G., and M.D. Welch. 2010. A nucleator arms race: cellular control of actin assembly. Nat Rev Mol Cell Biol. 11:237–251.

Cárcamo, C., E. Pardo, C. Oyanadel, M. Bravo-Zehnder, P. Bull, M. Cáceres, J. Martínez, L. Massardo, S. Jacobelli, A. González, and A. Soza. 2006. Galectin-8 binds specific beta1 integrins and induces polarized spreading highlighted by asymmetric lamellipodia in Jurkat T cells. Exp Cell Res. 312:374–386.

Case, L.B., and C.M. Waterman. 2011. Adhesive F-actin waves: a novel integrin-mediated adhesion complex coupled to ventral actin polymerization. PLoS One. 6:e26631.

Chhabra, E.S., and H.N. Higgs. 2007. The many faces of actin: matching assembly factors with cellular structures. Not Cell Biol. 9:1110–1121.

Cueni, L.N., and M. Detmar. 2009. Galectin-8 interacts with podoplanin and modulates lymphatic endothelial cell functions. Exp Cell Res. 315:1715–1723.

Devreotes, P., and A.R. Horwitz. 2015. Signaling networks that regulate cell migration. Cold Spring Harbor perspectives in biology. 7:a005959.

Diskin, S., Z. Cao, H. Leffler, and N. Panjwani. 2009. The role of integrin glycosylation in galectin-8-mediated trabecular meshwork cell adhesion and spreading. Glycobiology. 19:29–37.

Diskin, S., W.S. Chen, Z. Cao, S. Gyawali, H. Gong, A. Soza, A. González, and N. Panjwani. 2012. Galectin-8 promotes cytoskeletal rearrangement in trabecular meshwork cells through activation of Rho signaling. PLoS One. 7:e44400.

Elola, M.T., F. Ferragut, V.M. Cárdenas Delgado, L.G. Nugnes, L. Gentilini, D. Laderach, M.F. Troncoso, D. Compagno, C. Wolfenstein-Todel, and G.A. Rabinovich. 2014. Expression, localization and function of galectin-8, a tandem-repeat lectin, in human tumors. Histology and histopathology. 29:1093–1105.

Elola, M.T., C. Wolfenstein-Todel, M.F. Troncoso, G.R. Vasta, and G.A. Rabinovich. 2007. Galectins: matricellular glycan-binding proteins linking cell adhesion, migration, and survival. Cellular and molecular life sciences: CMLS. 64:1679–1700.

Fernández, M.M., F. Ferragut, V.M. Cárdenas Delgado, C. Bracalente, A.I. Bravo, A.J. Cagnoni, M. Nuñez, L.G. Morosi, H.R. Quinta, M.V. Espelt, M.F. Troncoso, C. Wolfenstein-Todel, K.V. Mariño, E.L. Malchiodi, G.A. Rabinovich, and M.T. Elola. 2016. Glycosylation-dependent binding of galectin-8 to activated leukocyte cell adhesion molecule (ALCAM/CD166) promotes its surface segregation on breast cancer cells. Biochimica et biophysica acta. 1860:2255–2268.

Ferragut, F., A.J. Cagnoni, L.L. Colombo, C. Sánchez Terrero, C. Wolfenstein-Todel, M.F. Troncoso, S.I. Vanzulli, G.A. Rabinovich, K.V. Mariño, and M.T. Elola. 2019. Dual knockdown of Galectin-8 and its glycosylated ligand, the activated leukocyte cell adhesion molecule (ALCAM/CD166), synergistically delays in vivo breast cancer growth. Biochimica et biophysica acta. Molecular cell research. 1866:1338–1352.

Flatau, G., E. Lemichez, M. Gauthier, P. Chardin, S. Paris, C. Fiorentini, and P. Boquet. 1997. Toxin-induced activation of the G protein p21 Rho by deamidation of glutamine. Nature. 387:729–733.

Gardel, M.L., I.C. Schneider, Y. Aratyn-Schaus, and C.M. Waterman. 2010. Mechanical integration of actin and adhesion dynamics in cell migration. Annual review of cell and developmental biology. 26:315–333.

Gauthier, N.C., T.A. Masters, and M.P. Sheetz. 2012. Mechanical feedback between membrane tension and dynamics. Trends Cell Biol. 22:527–535.

Geiger, B., J.P. Spatz, and A.D. Bershadsky. 2009. Environmental sensing through focal adhesions. Nat Rev Mol Cell Biol. 10:21–33.

Gentilini, L.D., F.M. Jaworski, C. Tiraboschi, I.G. Pérez, M.L. Kotler, A. Chauchereau, D.J. Laderach, and D. Compagno. 2017. Stable and high expression of Galectin-8 tightly controls metastatic progression of prostate cancer. Oncotarget. 8:44654–44668.

Guillaume-Gentil, O., E. Potthoff, D. Ossola, C.M. Franz, T. Zambelli, and J.A. Vorholt. 2014. Force-controlled manipulation of single cells: from AFM to FluidFM. Trends in biotechnology. 32:381–388.

Hadari, Y.R., R. Arbel-Goren, Y. Levy, A. Amsterdam, R. Alon, R. Zakut, and Y. Zick. 2000. Galectin-8 binding to integrins inhibits cell adhesion and induces apoptosis. Journal of cell science. 113 (Pt 13):2385–2397.

Hadari, Y.R., K. Paz, R. Dekel, T. Meštrović, D. Accili, and Y. Zick. 1995. Galectin-8: A NEW RAT LECTIN, RELATED TO GALECTIN-4. Journal of Biological Chemistry. 270:3447–3453.

He, J., and L.G. Baum. 2006. Galectin Interactions with Extracellular Matrix and Effects on Cellular Function. In Methods in enzymology. Vol. 417. Academic Press. 247–256.

Hu, S., K. Dasbiswas, Z. Guo, Y.-H. Tee, V. Thiagarajan, P. Hersen, T.-L. Chew, S.A. Safran, R. Zaidel-Bar, and A.D. Bershadsky. 2017. Long-range self-organization of cytoskeletal myosin II filament stacks. Nature Cell Biology. 19:133–141.

Humphrey, J.D., E.R. Dufresne, and M.A. Schwartz. 2014. Mechanotransduction and extracellular matrix homeostasis. Nat Rev Mol Cell Biol. 15:802–812.

Humphries, J.D., M.R. Chastney, J.A. Askari, and M.J. Humphries. 2019. Signal transduction via integrin adhesion complexes. Current opinion in cell biology. 56:14–21.

Innocenti, M. 2018. New insights into the formation and the function of lamellipodia and ruffles in mesenchymal cell migration. Cell Adh Migr. 12:401–416.

Jacquemet, G., H. Hamidi, and J. Ivaska. 2015. Filopodia in cell adhesion, 3D migration and cancer cell invasion. Current opinion in cell biology. 36:23–31.

Kassianidou, E., and S. Kumar. 2015. A biomechanical perspective on stress fiber structure and function. Biochimica et biophysica acta. 1853:3065–3074.

Kaufman, S.J., and M.L. Lawless. 1980. Thiodigalactoside Binding Lectin and Skeletal Myogenesis. Differentiation. 16:41–48.

Kengyel, A., W.A. Wolf, R.L. Chisholm, and J.R. Sellers. 2010. Nonmuscle myosin IIA with a GFP fused to the N-terminus of the regulatory light chain is regulated normally. Journal of muscle research and cell motility. 31:163–170.

Kühn, S., and M. Geyer. 2014. Formins as effector proteins of Rho GTPases. Small GTPases. 5:e29513.

Le Clainche, C., and M.F. Carlier. 2008. Regulation of actin assembly associated with protrusion and adhesion in cell migration. Physiological reviews. 88:489–513.

Levy, Y., R. Arbel-Goren, Y.R. Hadari, S. Eshhar, D. Ronen, E. Elhanany, B. Geiger, and Y. Zick. 2001. Galectin-8 Functions as a Matricellular Modulator of Cell Adhesion. Journal of Biological Chemistry. 276:31285–31295.

Levy, Y., S. Auslender, M. Eisenstein, R.R. Vidavski, D. Ronen, A.D. Bershadsky, and Y. Zick. 2006. It depends on the hinge: a structure-functional analysis of galectin-8, a tandem-repeat type lectin. Glycobiology. 16:463–476.

Medalia, O., and B. Geiger. 2010. Frontiers of microscopy-based research into cell-matrix adhesions. Current opinion in cell biology. 22:659–668.

Midwood, K.S., M. Chiquet, R.P. Tucker, and G. Orend. 2016. Tenascin-C at a glance. Journal of Cell Science. 129:4321–4327.

Midwood, K.S., and J.E. Schwarzbauer. 2002. Tenascin-C modulates matrix contraction via focal adhesion kinase-and Rho-mediated signaling pathways. Mol Biol Cell. 13:3601–3613.

Multhaupt, H.A.B., B. Leitinger, D. Gullberg, and J.R. Couchman. 2016. Extracellular matrix component signaling in cancer. Advanced Drug Delivery Reviews. 97:28–40.

Muncie, J.M., and V.M. Weaver. 2018. The Physical and Biochemical Properties of the Extracellular Matrix Regulate Cell Fate. Current topics in developmental biology. 130:1–37.

Naba, A., K.R. Clauser, H. Ding, C.A. Whittaker, S.A. Carr, and R.O. Hynes. 2016. The extracellular matrix: Tools and insights for the “omics” era. Matrix Biol. 49:10–24.

Nabi, I.R., J. Shankar, and J.W. Dennis. 2015. The galectin lattice at a glance. Journal of Cell Science. 128:2213–2219.

Newman, S.A., T. Glimm, and R. Bhat. 2018. The vertebrate limb: An evolving complex of self-organizing systems. Progress in biophysics and molecular biology. 137:12–24.

Nilufar, S., A.A. Morrow, J.M. Lee, and T.J. Perkins. 2013. FiloDetect: automatic detection of filopodia from fluorescence microscopy images. BMC Systems Biology. 7:66.

Otsu, N. 1979. A Threshold Selection Method from Gray-Level Histograms. IEEE Transactions on Systems, Man, and Cybernetics. 9:62–66.

Paran, Y., I. Lavelin, S. Naffar-Abu-Amara, S. Winograd-Katz, Y. Liron, B. Geiger, and Z. Kam. 2006. Development and application of automatic high-resolution light microscopy for cell-based screens. Methods in enzymology. 414:228–247.

Popa, S.J., S.E. Stewart, and K. Moreau. 2018. Unconventional secretion of annexins and galectins. Seminars in cell & developmental biology. 83:42–50.

Potthoff, E., O. Guillaume-Gentil, D. Ossola, J. Polesel-Maris, S. LeibundGut-Landmann, T. Zambelli, and J.A. Vorholt. 2012. Rapid and serial quantification of adhesion forces of yeast and Mammalian cells. PLoS One. 7:e527l2–e527l2.

Prager-Khoutorsky, M., A. Lichtenstein, R. Krishnan, K. Rajendran, A. Mayo, Z. Kam, B. Geiger, and A.D. Bershadsky. 2011. Fibroblast polarization is a matrix-rigidity-dependent process controlled by focal adhesion mechanosensing. Nat Cell Biol. 13:1457–1465.

Price, L.S., J. Leng, M.A. Schwartz, and G.M. Bokoch. 1998. Activation of Rac and Cdc42 by integrins mediates cell spreading. Mol Biol Cell. 9:1863–1871.

Ren, X.D., W.B. Kiosses, and M.A. Schwartz. 1999. Regulation of the small GTP-binding protein Rho by cell adhesion and the cytoskeleton. The EMBO journal. 18:578–585.

Resovi, A., D. Pinessi, G. Chiorino, and G. Taraboletti. 2014. Current understanding of the thrombospondin-1 interactome. Matrix Biology. 37:83–91.

Reticker-Flynn, N.E., D.F. Malta, M.M. Winslow, J.M. Lamar, M.J. Xu, G.H. Underhill, R.O. Hynes, T.E. Jacks, and S.N. Bhatia. 2012. A combinatorial extracellular matrix platform identifies cell-extracellular matrix interactions that correlate with metastasis. Nat Commun. 3:1122.

Romaniuk, M.A., G.A. Rabinovich, and M. Schattner. 2015. Galectins in the regulation of platelet biology. Methods in molecular biology (Clifton, N.J.). 1207:269–283.

Romaniuk, M.A., M.V. Tribulatti, V. Cattaneo, M.J. Lapponi, F.C. Molinas, O. Campetella, and M. Schattner. 2010. Human platelets express and are activated by galectin-8. The Biochemical journal. 432:535–547.

Sancho, A., I. Vandersmissen, S. Craps, A. Luttun, and J. Groll. 2017. A new strategy to measure intercellular adhesion forces in mature cell-cell contacts. Scientific Reports. 7:46152.

Schachtner, H., S.D. Calaminus, S.G. Thomas, and L.M. Machesky. 2013. Podosomes in adhesion, migration, mechanosensing and matrix remodeling. Cytoskeleton (Hoboken). 70:572–589.

Schell, M. J., C. Erneux, and R.F. Irvine. 2001. Inositol 1,4,5-trisphosphate 3-kinase A associates with Factin and dendritic spines via its N terminus. The Journal of biological chemistry. 276:37537–37546.

Schmidt, G., P. Sehr, M. Wilm, J. Selzer, M. Mann, and K. Aktories. 1997. Gln□63 of Rho is deamidated by Escherichia coli cytotoxic necrotizing factor-1. Nature. 387:725–729.

Shatz-Azoulay, H., Y. Vinik, R. Isaac, U. Kohler, S. Lev, and Y. Zick. 2020. The Animal Lectin Galectin-8 Promotes Cytokine Expression and Metastatic Tumor Growth in Mice. Scientific reports. 10:7375.

Small, J.V., K. Rottner, I. Kaverina, and K.I. Anderson. 1998. Assembling an actin cytoskeleton for cell attachment and movement. Biochimica et biophysica acta. 1404:271–281.

Staus, D.P., J.M. Taylor, and C.P. Mack. 2011. Enhancement of mDia2 activity by Rho-kinase-dependent phosphorylation of the diaphanous autoregulatory domain. The Biochemical journal. 439:57–65.

Steffen, A., S.A. Koestler, and K. Rottner. 2014. Requirements for and consequences of Rac-dependent protrusion. European journal of cell biology. 93:184–193.

Swaney, K.F., and R. Li. 2016. Function and regulation of the Arp2/3 complex during cell migration in diverse environments. Current opinion in cell biology. 42:63–72.

Tojkander, S., G. Gateva, and P. Lappalainen. 2012. Actin stress fibers--assembly, dynamics and biological roles. J Cell Sci. 125:1855–1864.

Troncoso, M.F., F. Ferragut, M.L. Bacigalupo, V.M. Cárdenas Delgado, L.G. Nugnes, L. Gentilini, D. Laderach, C. Wolfenstein-Todel, D. Compagno, G.A. Rabinovich, and M.T. Elola. 2014. Galectin-8: a matricellular lectin with key roles in angiogenesis. Glycobiology. 24:907–914.

Vinik, Y., H. Shatz-Azoulay, A. Vivanti, N. Hever, Y. Levy, R. Karmona, V. Brumfeld, S. Baraghithy, M. Attar-Lamdar, S. Boura-Halfon, I. Bab, and Y. Zick. 2015. The mammalian lectin galectin-8 induces RANKL expression, osteoclastogenesis, and bone mass reduction in mice. eLife. 4:e05914.

Vinik, Y., H. Shatz-Azoulay, and Y. Zick. 2018. Molecular Mechanisms Underlying the Role of Galectin-8 as a Regulator of Cancer Growth and Metastasis. Trends in Glycoscience and Glycotechnology. 30:SEII9–SEI28.

Walko, G., M.J. Castañón, and G. Wiche. 2015. Molecular architecture and function of the hemidesmosome. Cell Tissue Res. 360:529–544.

Wenk, M.B., K.S. Midwood, and J.E. Schwarzbauer. 2000. Tenascin-C suppresses Rho activation. J Cell Biol. 150:913–920.

Wilkinson, S., H.F. Paterson, and C.J. Marshall. 2005. Cdc42–MRCK and Rho-ROCK signalling cooperate in myosin phosphorylation and cell invasion. Nature Cell Biology. 7:255–261.

Wolfenson, H., Y.I. Henis, B. Geiger, and A.D. Bershadsky. 2009. The heel and toe of the cell’s foot: a multifaceted approach for understanding the structure and dynamics of focal adhesions. Cell Motil Cytoskeleton. 66:1017–1029.

Wolfenson, H., T. Iskratsch, and M.P. Sheetz. 2014. Early events in cell spreading as a model for quantitative analysis of biomechanical events. Biophysical journal. 107:2508–2514.

Xia, S., and P. Kanchanawong. 2017. Nanoscale mechanobiology of cell adhesions. Seminars in cell & developmental biology. 71:53–67.

York, A.G., P. Chandris, D.D. Nogare, J. Head, P. Wawrzusin, R.S. Fischer, A. Chitnis, and H. Shroff. 2013. Instant super-resolution imaging in live cells and embryos via analog image processing. Nature methods. 10:1122–1126.

Zaidel-Bar, R., R. Milo, Z. Kam, and B. Geiger. 2007. A paxillin tyrosine phosphorylation switch regulates the assembly and form of cell-matrix adhesions. Journal of Cell Science. 120:137–148.

Zhao, Z., and E. Manser. 2015. Myotonic dystrophy kinase-related Cdc42-binding kinases (MRCK), the ROCK-like effectors of Cdc42 and Rac1. Small GTPases. 6:81–88.

Zick, Y., M. Eisenstein, R.A. Goren, Y.R. Hadari, Y. Levy, and D. Ronen. 2002. Role of galectin-8 as a modulator of cell adhesion and cell growth. Glycoconjugate Journal. 19:517–526.

