## Supplementary Materials for "Differential cellular responses to adhesive interactions with galectin-8 and fibronectin coated substrates"

**Key words:**

Mechanobiology Institute

National University of Singapore

Singapore

Prof. Benjamin Geiger

Department of Immunology

Weizmann Institute of Science

Rehovot 7610001, Israel

T: +972-8-934 3910

M: +972-52-348 8848

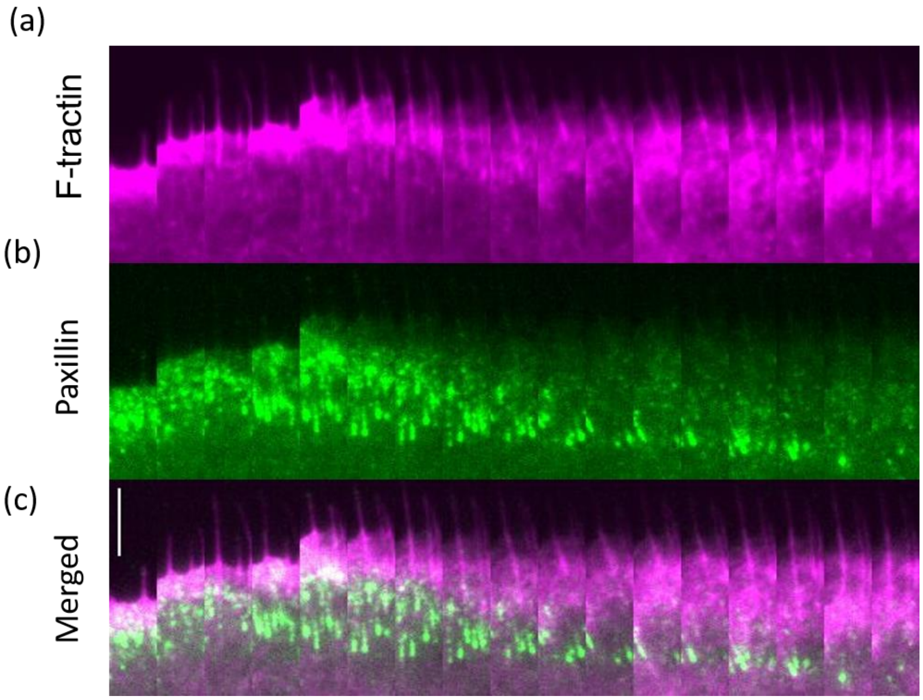


**Supplementary Figure 1: Kymograph showing the dynamics of lamellipodia and paxillin clusters in cells plated on galectin-8**. The cell was labeled with tdTomato-F-tractin and YFP-paxillin, and imaged 20 minutes after plating on a galectin-8 coated substrate. The rectangular strip perpendicular to the cell's leading edge was filmed every 2 minutes. Note formation of numerous paxillin clusters associated with actin-enriched lamellipodia, which moved centripetally and eventually disappeared. Scale bar: 5 µm.

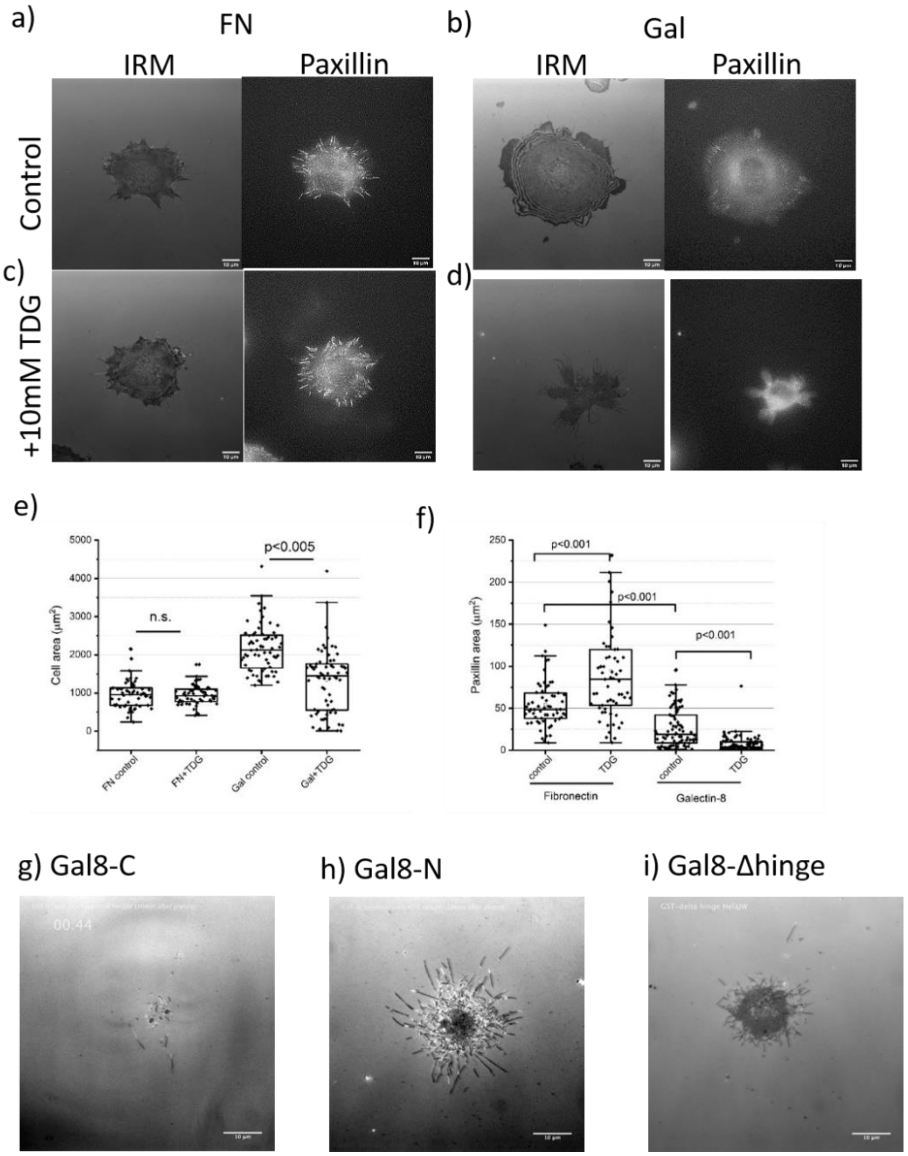


**Supplementary Figure 2**: Thiodigalactoside (TDG) addition inhibits cell spreading and adhesion formation on galectin-8 but not on fibronectin. (a, b) IRM and YFP-paxillin images of cells on fibronectin (a, c) and on galectin-8 (b, d) at 0.5 hour of spreading without (a, b) or with 10mM TDG (c, d). (e, f) Quantification of projected cell area (e) and total paxillin area per cell (f) upon spreading on fibronectin and galectin-8 in the presence or absence of TDG. (g-h) Cell spreading on different galectin-8 mutants.

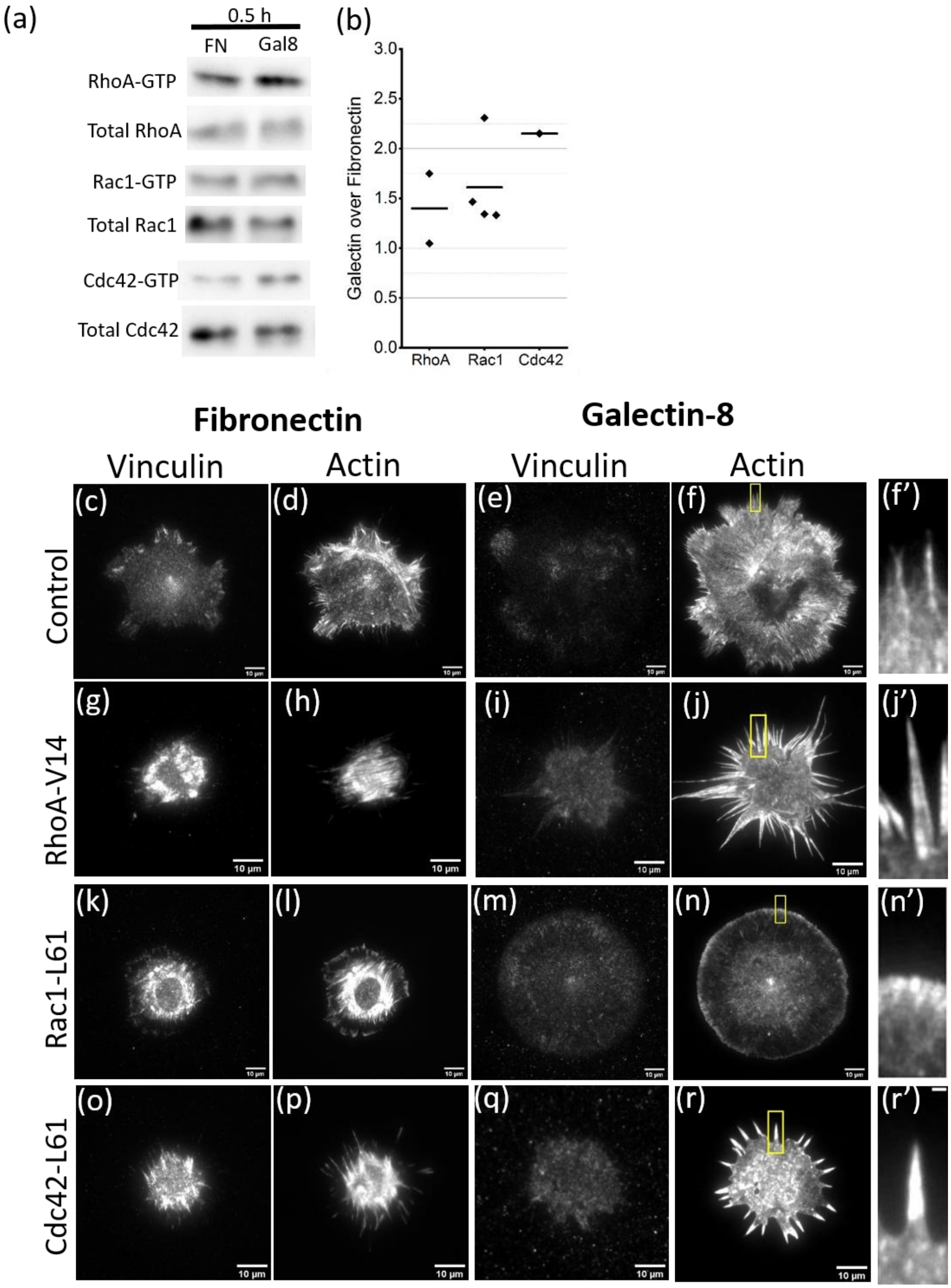

(a)
0.5 h
FN
Gal8
RhoA-GTP
Total RhoA
Rac1-GTP
Total Rac1
Cdc42-GTP
Total Cdc42
(b)
Galectin over Fibronectin
3.0
2.5
2.0
1.5
1.0
0.5
0.0
RhoA
Rac1
Cdc42
Fibronectin
Galectin-8
Vinculin
Actin
Vinculin
Actin
Control
RhoA-V14
Rac1-L61
Cdc42-L61
(c)
(d)
(e)
(f)
(f')
(g)
(h)
(i)
(j)
(j')
(k)
(l)
(m)
(n)
(n')
(o)
(p)
(q)
(r)
(r')
10 μm

**Supplementary Figure 3: Effects of small Rho GTPases on cell spreading on fibronectin- and galectin-8 coated substrates.** (a) Pull-down assay of small GTPase activity. Western blot showing active and total RhoA, Rac1, Cdc42 levels in cells plated on fibronectin and galectin-8 for 30 minutes. (b) Ratios of RhoA, Rac1, and Cdc42 activity 30 minutes following plating on galectin-8, compared to those following plating on fibronectin. Each dot corresponds to independent experiments. Mean values are indicated by horizontal lines. (c-r) Paxillin-positive adhesions (c,e,i,k,m,o,q) and actin structures (d,f,h,j,l,n,p) in control cells (c-f) and cells expressing constitutively active RhoA-V14 (g-j), Rac1-L61(k-n), and Cdc42-L61(o-r), 2 hours following plating on fibronectin-(c,d,g,h,k,l,o,p) or galectin-8(e,f,i,j,m,n,q,r) coated substrates. Cells were fixed and stained with paxillin antibody and TRITC-phalloidin to visualize F-actin. Scale bars: 10 µm. Boxed areas in f, j, n, r at high magnification are shown in f', j', n', r', respectively. Scale bar shown in r': 2 µm.

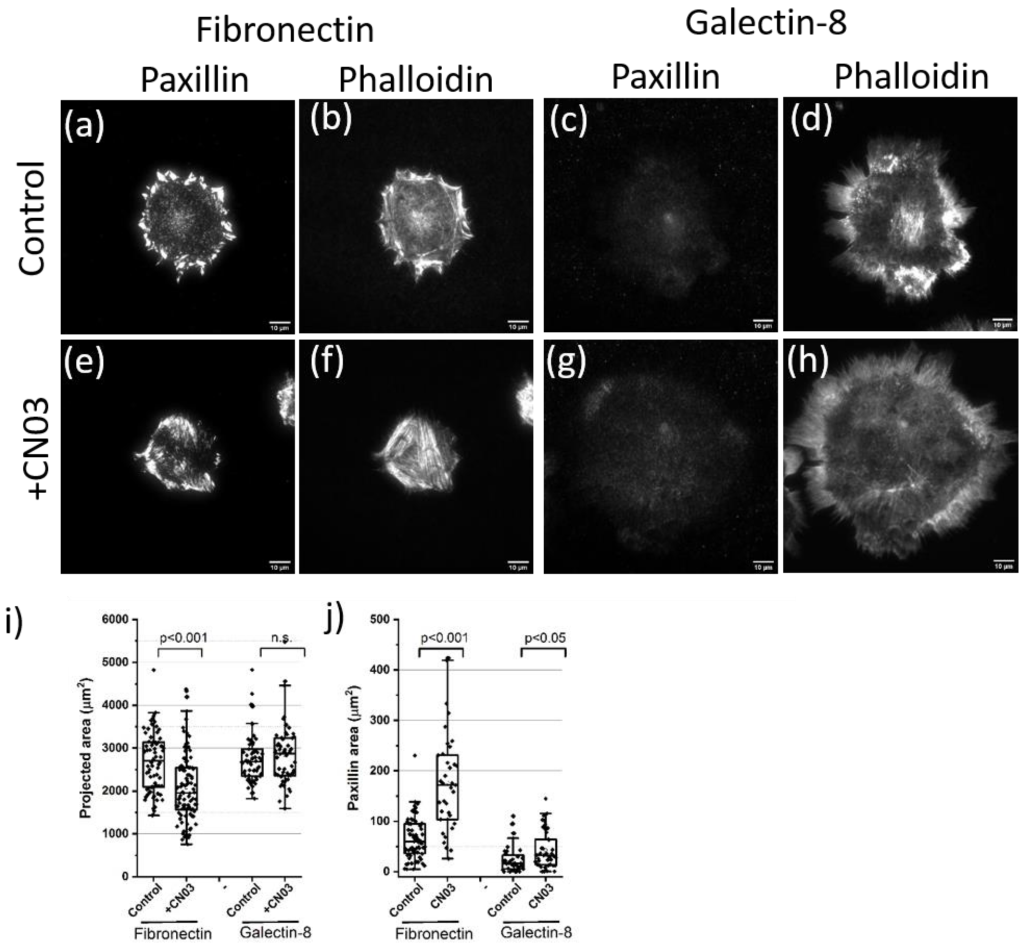


**Supplementary Figure 4: Effects of pharmacological RhoA activation on formation of paxillin-positive adhesions and actin cytoskeleton structures in cells spreading on fibronectin and galectin-8 substrates.** Cells were plated on fibronectin- (a,b,e,f) or galectin-8 (c,d,g,h) coated substrates in the absence (a,b,c,d) or presence (e,f,g,h) of 1 µM RhoA activator CN03, and fixed 3 hours following plating. Paxillin adhesions were visualized by antibody to paxillin (a,e,c,g), and F-actin, by TRITC-phalloidin staining (b,d,f,h). Scale bar: 10 µm. (i,j) Quantification of projected cell area (i) and total area of paxillin clusters per cell (j) in control cells, and in cells treated with 1 µM CN03. Morphometric measurements and presentation of results were performed as described in the legend to Figure 6.

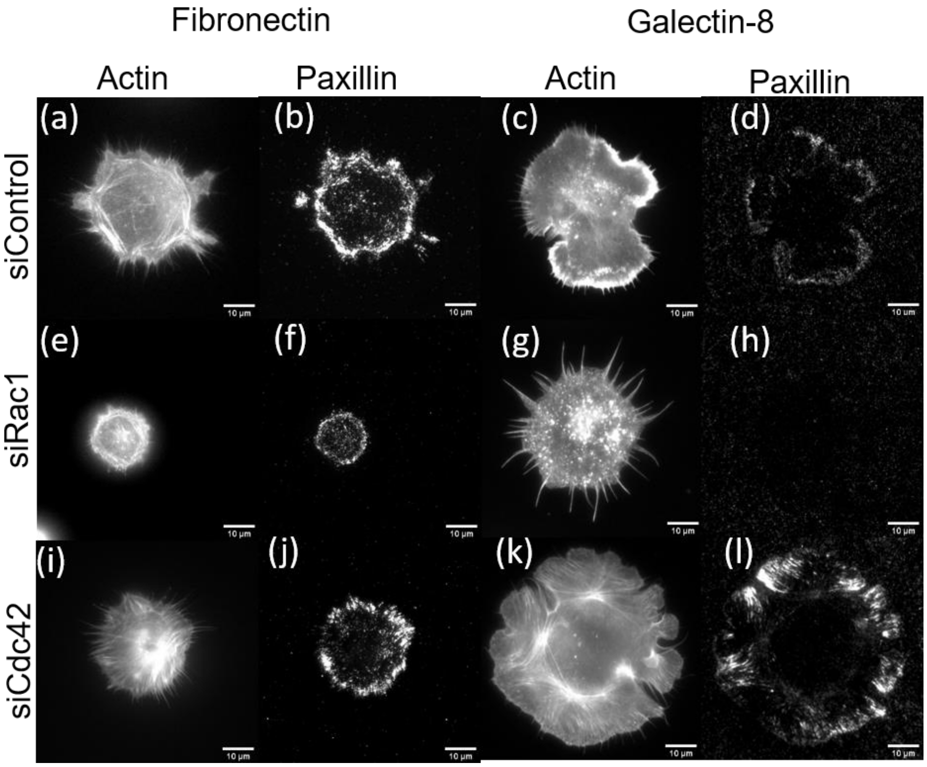


**Supplementary Figure 5: Effects of depletion of small Rho GTPases on cell spreading on fibronectin and galectin-8 coated substrates**. Cells transfected with control siRNA (a-d), siRNA against Rac1 (e-h) and Cdc42 (i-l) were plated on fibronectin (a,b,e,f,i,j) and galectin-8 (c,d,g,h,k,l) coated substrates, and fixed 30 minutes following plating. Cells were stained with TRITC-phalloidin to visualize F-actin (a,e,i,c,g,k) and with antibody to paxillin (b,f,j,d,h,l). Scale bars: 10 µm.

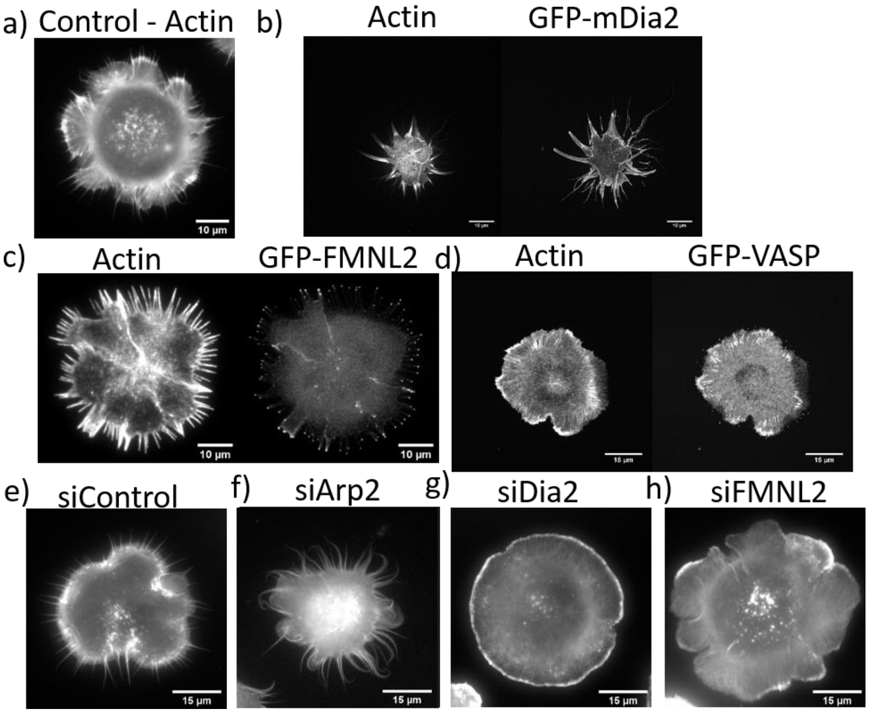


**Supplementary Figure 6: Effects of downstream effectors of Rho family G-proteins on cell spreading on galectin-8 coated substrates**. Control cells (a), cells expressing the constitutively active formins EGFP-mDia2 (b) and EGFP-FMNL2 (c), cells over-expressing actin polymerization activator EGFP-VASP (d), as well as cells expressing control siRNA (e) and siRNAs to Arp2 (f), mDia2 (g), and FMNL2(h) were plated on a galectin-8 coated substrate, fixed 20 minutes following plating, and stained with TRITC-phalloidin to visualize F-actin. In (b), (c), and (d), actin images (left) and images showing the localization of EGFP-mDia2, EGFP-FMNL2, EGFP-VASP in the same cells (right) are shown. Morphometric measurements and presentation of results were performed as described in the legend to Figure 6.

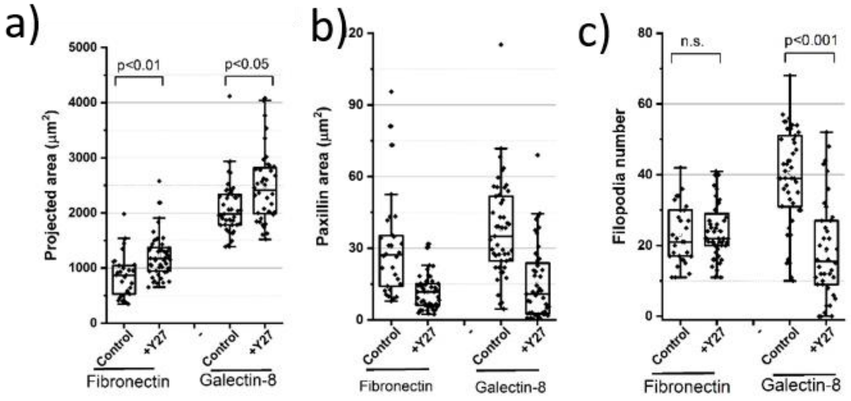


**Supplementary Figure 7: Effects of inhibition of Rho kinase (ROCK) on cell spreading on fibronectin and galectin-8 substrates.** Quantification of projected cell areas (a), total areas of paxillin clusters per cell (b), and numbers of filopodia (c) in cells plated on fibronectin and galectin-8 substrates in the absence (control) and the presence of 100 μM of Y27632. Morphometric measurements and presentation of results were performed as described in the legend to Figure 6.

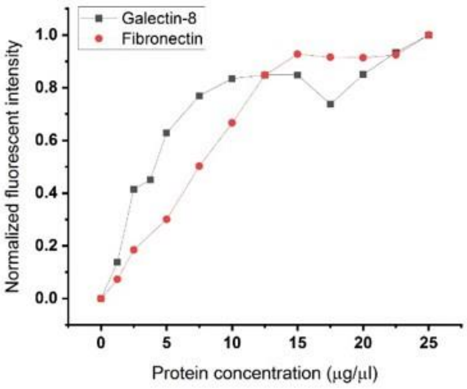


**Supplementary Figure 8: Protein absorption on glass bottomed petri-dish.** Note that protein absorption on the surface reaches plateau at the concentration of 15 μg/μl for both galectin-8 and fibronectin.

**Video Captions**

Movie 1: **Spreading of cells on fibronectin coated substrates.** Representative time-lapse series showing cells spreading fibronectin-coated substrate, imaged by interference reflection microscopy (IRM) (left) and differential interference contrast microscopy (DIC) (right). The time interval is 5 seconds. Scale bar: 15 µm. Timestamp: mm:ss, The display rate: 15 frames $sec^{-1}$.

Movie 2: **Spreading of cells on galectin-8 coated substrates.** Representative time-lapse series showing cells spreading fibronectin-coated substrate, imaged by interference reflection microscopy (IRM) (left) and differential interference contrast microscopy (DIC) (right). The time interval is 5 seconds. Scale bar: 15 µm. Timestamp: mm:ss, The display rate: 15 frames $sec^{-1}$.

Movie 3: **Spreading of cells on galectin-8 coated substrates.** Representative time-lapse series showing cells spreading fibronectin-coated substrate, imaged by interference reflection microscopy (IRM). The time interval is 2 seconds. Scale bar: 15 µm. Timestamp: mm:ss, The display rate: 15 frames $sec^{-1}$.

Movie 4: **Actin and myosin II dynamics during cell spreading on fibronectin.** Representative time-lapse series showing cells spreading fibronectin-coated substrate, imaged by SIM. Cells were labeled with tdTomato-F-tractin (actin) and GFP myosin II regulatory light chain (MRLC). The time interval is 9.5 seconds. Scale bar: 10 µm. Timestamp: mm:ss, The display rate: 15 frames $sec^{-1}$.

Movie 5: **Actin and myosin II dynamics during cell spreading on galectin-8.** Representative time-lapse series showing cells spreading galectin-8-coated substrate, imaged by SIM. Cells were labeled with tdTomato-F-tractin (actin) and GFP myosin II regulatory light chain (MRLC). The time interval is 10 seconds. Scale bar: 10 µm. Timestamp: mm:ss, The display rate: 15 frames $sec^{-1}$.

Movie 6: **Paxillin dynamics during cell spreading on fibronectin.** Representative time-lapse series showing cells spreading fibronectin-coated substrate, imaged by TIRF. Cells were labeled with YFP-paxillin. . The time interval is 30 seconds. Scale bar: 10 µm. Timestamp: mm:ss, Playback rate: 15 frames $sec^{-1}$.

Movie 7: **Paxillin dynamics during cell spreading on galectin-8.** Representative time-lapse series showing cells spreading galectin-8-coated substrate, imaged by SIM. Cells were labeled with YFP-paxillin. The time interval is 30 seconds. Scale bar: 10 µm. Timestamp: mm:ss, Playback rate: 15 frames $sec^{-1}$.

Movie 8: **Spreading of cells expressing constitutively active RhoA on galectin-8 coated substrates.** Representative time-lapse series showing cells spreading galectin-8-coated substrate, imaged by interference reflection microscopy (IRM). The time interval is 5 seconds. Scale bar: 10 µm. Timestamp: mm:ss, Playback rate: 15 frames $sec^{-1}$.

Movie 9: **Spreading of cells expressing constitutively active Rac1 on galectin-8 coated substrates.** Representative time-lapse series showing cells spreading galectin-8-coated substrate, imaged by interference reflection microscopy (IRM). The time interval is 5 seconds. Scale bar: 10 µm. Timestamp: mm:ss, Playback rate: 15 frames $sec^{-1}$.
